# Population genomics reveals fine-scale three-dimensional structure within two sympatric *Sebastes* species in the Northwest Atlantic

**DOI:** 10.64898/2025.12.01.686997

**Authors:** Audrey Bourret, Hugues P. Benoît, Caroline Senay, Geneviève J. Parent

## Abstract

An ideal fishery stock assessment requires a comprehensive understanding of population structure across landscapes. In the Gulf of St. Lawrence and the Laurentian Channel (GSL-LC), massive *Sebastes* recruitments occurred in early 2010s, nearly 30 years after the last strong cohort. This recruitment had resulted in abundant *Sebastes mentella*, but also involved the morphologically nearly indistinguishable *S. fasciatus*. Both species show multiple populations, but spatially explicit information at a management-relevant scale is lacking to support robust scientific advices for sustainable fisheries. Temporal variation in the *Sebastes* recruitment also remains poorly characterized, and it is unknown if the recruitment is synchronized at the species or population level. This study aimed to 1) characterize the current fine-scale genomic structure of *S. mentella* and *S. fasciatus* in the GSL-LC, 2) compare the genetic composition of different cohorts, and 3) evaluate relationship between genomic structure and two key factors in redfish management, depth, and management units. Our genomic datasets (> 16,000 SNPs, N = 2,248 redfish) revealed substructure within the previously identified *S. mentella GSL* ecotype and five *S. fasciatus* populations within the GSL-LC. While all genetic groups were represented in the recent cohort samples, our results suggested unequal contributions of *S. fasciatus* populations to massive recruitment events. The spatial distribution of genetic groups within both species revealed a three-dimensional structure tied to management units and depth. Our findings underscore the importance of revising management measures to incorporate population structure thereby reducing the risk of overexploiting smaller populations, particularly *S. fasciatus*, and promoting sustainable fisheries.

## Introduction

Best practices in marine resource management consider the underlying biological units (Cadrin 2020). Adequation between management and biological units reduce risks of overexploitation of a stock, enhance recovery of depleted stocks, improved sustainable yields, and protect the genetic and demographic diversity (Stephenson 1999; Cadrin and Secor 2009; Pinsky and Palumbi 2014; Hilborn et al. 2020). However, management area boundaries may not align with those of biological units for numerous reasons, including limited data or insufficient resolution of assessment methods, which often result in poorly understood biological units (Reiss et al. 2009; Kerr et al. 2017; Cadrin 2020). Genomic approaches provide powerful tools for investigating fine-scale genomic structure, identifying biological units such as species and populations, and characterizing their temporal stability.

In the Northwest Atlantic, two species mainly support redfish fisheries, the Deep water redfish, *Sebastes mentella,* and the Acadian redfish, *S. fasciatus*. A third species, *S. norvegicus*, is only occasionally encountered in this region and is more typical of the Northeast Atlantic. Similarly to other *Sebastes*, they are ovoviviparous (i.e., internal fertilization and release of larvae) and long lived species (>65 years). *S. mentella* is generally associated with northern and deeper habitats, while *S. fasciatus* tends to occupy southern and shallower areas. Nevertheless, their distributions overlap broadly, from the Laurentian Chanel to the Labrador Sea, including the Gulf of St. Lawrence (GSL; Fig. 1). These two species are highly morphologically similar and cannot be reliably discriminated macroscopically in the field (Gascon 2003; Senay et al. 2022). In areas of sympatry, where both species co-occur, accurate identification remains particularly problematic, and consequently both species are sometimes managed as a single stock within a management unit (DFO 2020, 2023).

**Figure 1.**
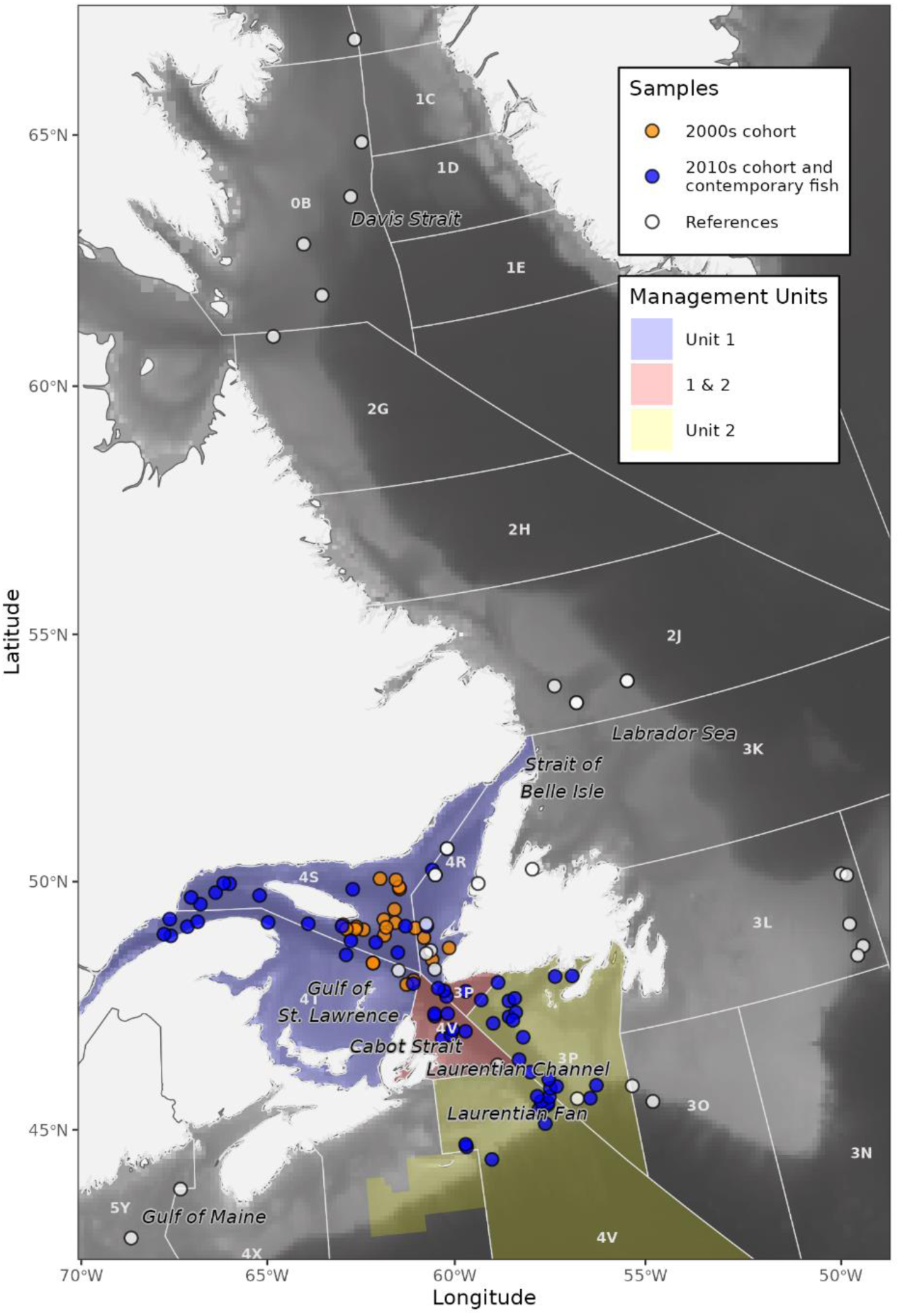
: Distribution of *Sebastes* samples for the 2000s cohorts, 2010s cohort and contemporary fish, and the reference datasets in the Northwest Atlantic. NAFO divisions (white line) and management units (colored areas) and important seascape features are presented.

Although species discrimination is morphologically difficult, genetic markers discriminate easily *S. mentella* and *S. fasciatus*. Beyond interspecific differentiation, significant intraspecific genetic structure has been demonstrated and should be considered for management based on biological units (Valentin et al. 2014; Saha et al. 2021; Benestan et al. 2021; Jansson et al. 2025). Within *S. mentella*, three ecotypes were described in the Northwest Atlantic, i.e., “*Deep*”, “*Shallow*”, and “*GSL*” ecotypes. The latter is specific to the Gulf of St. Lawrence, including the extension of the Laurentian Channel to the Atlantic Ocean (GSL-LC; Benestan et al. 2021; Fig. 1). For *S. fasciatus*, at least five populations were observed, but their exact number and distribution are not well defined yet (Benestan et al. 2021). The GSL-LC is also recognized as an area where both species display genetic signals of introgression, i.e., hybridization and then backcross with parental species (Roques et al. 2001; Valentin et al. 2014; Benestan et al. 2021). The complex structure and limited knowledge on ecotypes and populations continue to challenge the optimal management of *Sebastes* species in the Northwest Atlantic in general, and the GSL-LC in particular.

Since the mid-1990s, the GSL-LC has been divided into two management units, Unit 1 and Unit 2, to manage the co-occurring *S. mentella* and *S. fasciatus* (Fig. 1). These boundaries were set based on biological and spatiotemporal considerations. Across the year, Unit 1 includes NAFO Divisions 4RST whereas Unit 2 includes NAFO Subdivisions 3Ps4Vs4Wfgj (Senay et al. 2023).The NAFO Subdivisions 3Pn4Vn switch between Unit 1 (January to May) and Unit 2 (June to December) to reflect suspected seasonal redfish movements through the Cabot Strait, and referred hereafter as the shared area Unit 1&2 (Fig. 1). Within these spatio-temporal management boundaries, bottom-depth plays a key role in redfish ecology and management in the GSL-LC. Notably, *S. mentella* is predominantly associated with deeper habitats (>250 m), whereas *S. fasciatus* is more commonly found in shallower zones (100-300m; Senay et al. 2023). Since depth is a readily measurable and consistent environmental variable, current management measures include a minimum fishing depth of 183 m all year round.

Concerns were raised regarding the validity of these management units, owing to a lack of genetic structure using microsatellites (Valentin et al. 2014). In contrast, Benestan et al. (2021) showed some genetic structure using more powerful markers (thousands of single-nucleotide polymorphisms, SNPs), but the limited sampling within the GSL-LC prevented a conclusive assessment of the management unit’s relevance. Thus, there remains a need for a finer-scale study to assess the relevance of current management units.

Episodic recruitment adds yet another layer of complexity to the management of redfish. Long-lived *Sebastes*, including *S. mentella* and *S. fasciatus*, are characterized by highly variable recruitment, with large year classes observed at irregular intervals (Gascon 2003; Licandeo et al. 2020; Cadigan et al. 2022). In years 2011, 2012 and 2013, strong recruitment of redfish was observed in the GSL-LC (hereafter, the 2010s cohorts). These cohorts comprised mainly *S. mentella* although *S. fasciatus* also contributed to a smaller extent (approx. 10%, Brassard et al. 2017). The previous important cohort in the Unit 1 dates back to the 1980s, and also mainly comprised *S. mentella* (Valentin et al. 2015). Some large cohorts of *S. fasciatus* were observed in the GSL-LC (e.g., 1988, 2003), but these disappeared at ca. 5-6 years of age for reasons that remain unclear. Moreover, it is uncertain whether recruitment patterns are consistent across the different populations (Valentin et al. 2015). Sustainable management of redfish populations should therefore include a better understanding of recruitment dynamics and their links to genetic diversity.

In this study, we aimed to address key knowledge gaps and our objectives were (1) to characterize the current genomic structure of *S. mentella* and *S. fasciatus* in the GSL-LC, (2) to contrast the current genomic composition of the 2010s cohorts to that of previous cohorts, and (3) to evaluate the association of genomic structure with depth and management units (Unit 1, Unit2, and the shared Unit 1&2), the two main axes of redfish management in the region.

## Methods

### Sampling

For this study, we collected *Sebastes* specimens representative of (1) the 2000s cohort, (2) the 2010s cohorts and contemporary fish in the GSL-LC, and (3) *Sebastes* diversity across the Northwest Atlantic, hereafter called “References” (Table 1, Fig. 1). All specimens were sampled for tissues (fin or muscle), which were preserved in 95% ethanol until genetic analysis. In addition, fork length was measured and sex was recorded when possible. Fish within the GSL-LC were attributed to a management unit, i.e., Unit 1 (NAFO 4RST), Unit 2 (NAFO 3Ps4Vs4Wfgj), or the shared area Unit 1&2 (NAFO 3Pn4Vn; Fig. 1).

**Table 1:**
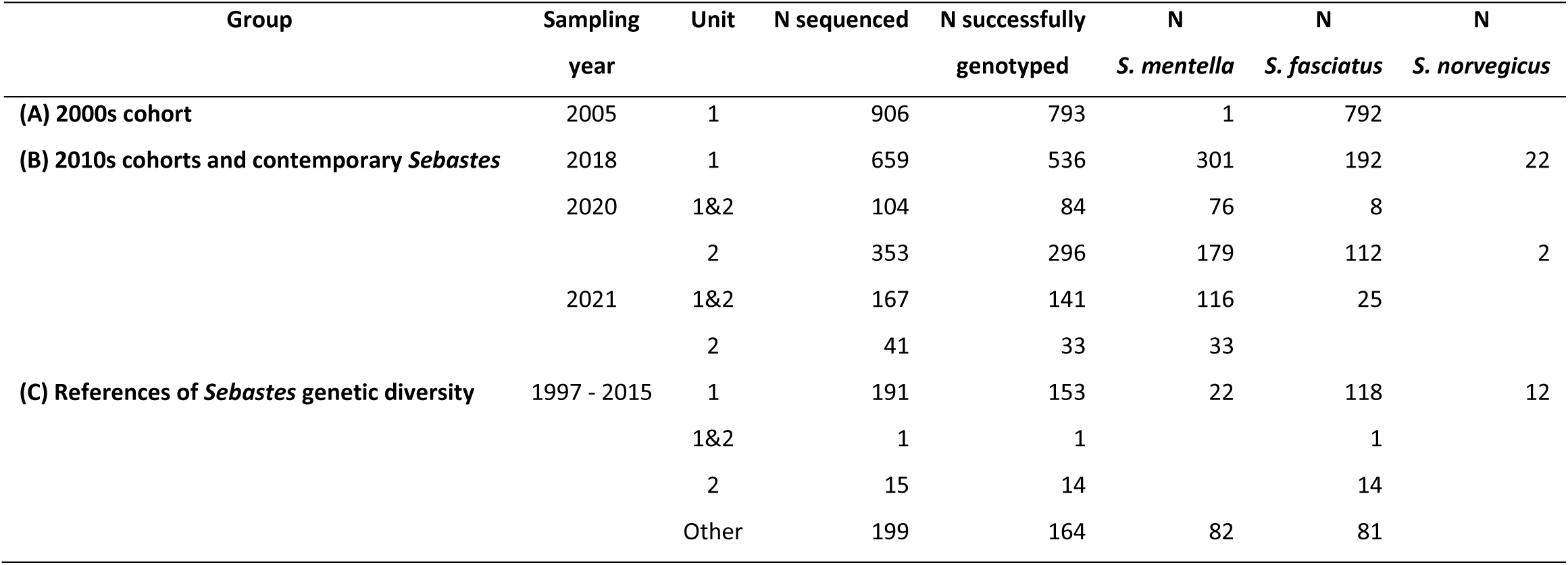
Details on the number of *Sebastes* sequenced and genotyped for the (A) 2000s cohort and (B) 2010s cohort and contemporary fish in the Gulf of St. Lawrence and Laurentian Chenal (Unit 1, 1&2, 2), and the (C) reference of *Sebastes* genetic. Individuals were assigned to *S. mentella, S. fasciatus* or *S. norvegicus* using the *Sebastes* spp. dataset, except for those identified as introgressed which were not assigned. See Fig. 1 for a map of all sampling locations.

### 2000s cohort

The 2000s cohort *Sebastes* were collected during the annual Department of Fisheries and Oceans Canada (DFO) ecosystemic survey in NAFO Divisions 4RST in August 2005 (Bourdages et al. 2007). During this survey, 21 to 99 individuals were randomly selected at 22 stations for a total of 975 specimens.

Sampled individuals ranged in length from 53 to 105 mm (µ: 85.9 ± 7.6 [standard deviation, SD] mm), and were suspected to be two years old.

### 2010s cohorts and contemporary *Sebastes*

The 2010s cohorts and contemporary *Sebastes* were collected from three sources. In 2018, three to 50 individuals were randomly selected across fish > 220 mm at 24 stations from the annual DFO ecosystemic survey for a total of 906 individuals. Their size varied from 124 to 468 mm (µ: 269.2 ± 58.4 mm). A summer redfish survey led by the industry, the Atlantic Groundfish Council, has been carried in NAFO Subdivisions 3Pn, 3Ps, 4Vn, and 4Vs in many years since 1997 (Kulka and Atkinson 2016). During their 2020 survey, between one and 39 individuals were randomly selected within 39 stations for a total of 694 specimens, which ranged in length from 102 to 425 mm (µ: 250.3 ± 37.0 mm). In order to improve the spatial coverage in NAFO Subdivisions 3Ps and 4Vn, between 50 and 52 individuals were randomly selected during five commercial fishing events in 2021, for a total of 253 specimens, which ranged in length from 174 to 395 mm (µ: 266.0 ± 26.7 mm).

#### References

To represent the previously described *Sebastes* genetic diversity within the Northwest Atlantic, 394 redfish sampled between 2001 and 2015 from sites ranging from the Gulf of Maine to the Davis Strait were included as “References” for the three *S. mentella* ecotypes and the multiple *S. fasciatus* populations (Benestan et al. 2021; Table 1, Fig. 1). Moreover, 35 *S. norvegicus* were added to characterize potential introgression with this *Sebastes* species that can occasionally be encountered in the GSL-LC (Table 1). These references of *Sebastes* genetic diversity ensure that the genetic variability observed in a new dataset is contrasted to the global known genetic variation across the distributional range.

### Genotyping

#### DNA extraction, library preparation, and sequencing

DNA was extracted with DNeasy Blood and Tissus or DNeasy 96 (Qiagen) and quantified on a Synergy LX (BioTek) using PicoGreen to select only high quality DNA extracts. Double digest restriction-site-associated DNA (ddRAD; *PstI* and *MspI* enzymes) libraries were prepared by the Plateforme d’Analyse Génomique (IBIS, Université Laval) using 20 ng of DNA per sample following Poland et al. (2012). A total of 2,660 individuals and 28 duplicates were pooled in 28 libraries of 96 samples and were sequenced on two lanes of NovaSeq X PE150 at Genome Quebec, with 10% PhiX.

#### SNPs datasets

Genotyping was performed using STACKS modules (v.2.55, Catchen et al. 2013; Rochette et al. 2019; Table 2). Read quality was checked using FastQC (Andrews 2010) and multiQC (Ewels et al. 2016), and Illumina adaptors were removed with Trimmomatic (Bolger et al. 2014). Reads were demultiplexed with *process_radtag* module, with a truncation at 120 pb, then aligned to the chromosome-level *Sebastes aleutianus* genome (Kolora et al. 2021) using BWA-MEM (Li and Durbin 2009) with default parameters. We used the *gstacks* module to create a SNPs catalog and to genotype all good quality samples (Table 2). Different filtration steps were performed to keep only SNPs shared by > 75% individuals (*population* modules, r = 0.75, MAF 0.01), low missingness SNPs (<10%) and individuals (<30%), and to remove SNPs with observed heterozygosity >0.6 and with too low (<10x) or high depth (>51x), and individuals with <5x coverage and high kinship coefficient (Φ>0.3, (Manichaikul et al. 2010) algorithm implemented in VCFtools, (Danecek et al. 2011). Three different datasets were obtained to answer inter and intra- specific questions, with MAF 1 or 5%, and one SNP by loci (see Table 2 for detailed filtration steps and SNPs datasets).

**Table 2:**
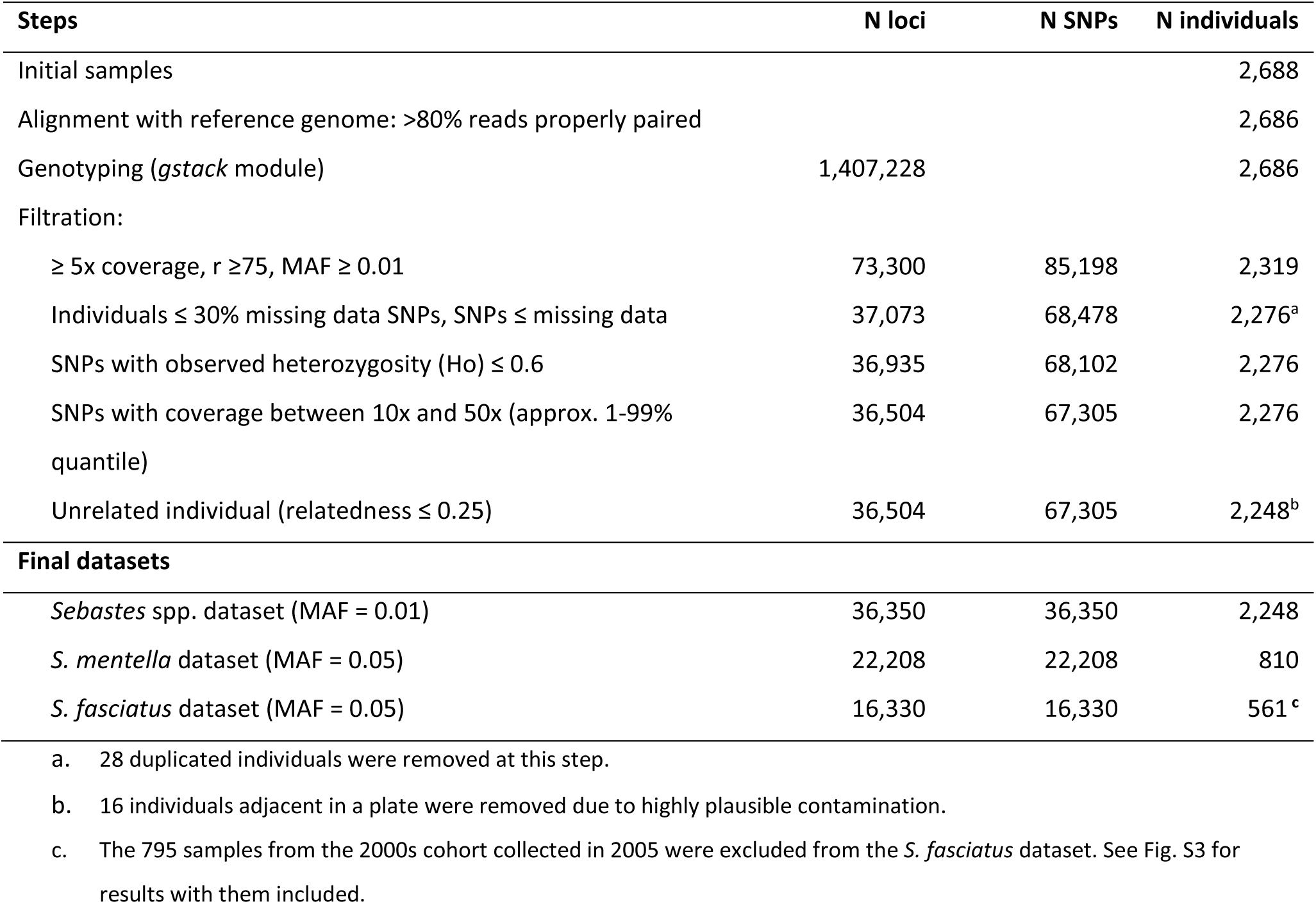
Number of loci, SNPs and individuals at each pipeline step and for the three final datasets. For the *S. mentella* and *S. fasciatus* species specific datasets, individuals identified as introgressed and SNPs within sex-linked genomic regions were removed. Final datasets kept only 1 SNP by loci.

| Steps | N loci | N SNPs | N individuals |
| --- | --- | --- | --- |
| Initial samples |  |  | 2,688 |
| Alignment with reference genome: >80% reads properly paired |  |  | 2,686 |
| Genotyping (*gstack* module) | 1,407,228 |  | 2,686 |
| Filtration: |  |  |  |
| $\geq 5x$ coverage, $r \geq 75$, $MAF \geq 0.01$ | 73,300 | 85,198 | 2,319 |
| Individuals $\leq 30\%$ missing data SNPs, SNPs ≤ missing data | 37,073 | 68,478 | 2,276<sup>a</sup> |
| SNPs with observed heterozygosity ($H_o$) $\leq 0.6$ | 36,935 | 68,102 | 2,276 |
| SNPs with coverage between 10x and 50x (approx. 1-99% quantile) | 36,504 | 67,305 | 2,276 |
| Unrelated individual (relatedness $\leq 0.25$) | 36,504 | 67,305 | 2,248<sup>b</sup> |
| **Final datasets** |  |  |  |
| *Sebastes* spp. dataset ($MAF = 0.01$) | 36,350 | 36,350 | 2,248 |
| *S. mentella* dataset ($MAF = 0.05$) | 22,208 | 22,208 | 810 |
| *S. fasciatus* dataset ($MAF = 0.05$) | 16,330 | 16,330 | 561<sup>c</sup> |

a. 28 duplicated individuals were removed at this step.
b. 16 individuals adjacent in a plate were removed due to highly plausible contamination.
c. The 795 samples from the 2000s cohort collected in 2005 were excluded from the *S. fasciatus* dataset. See Fig. S3 for results with them included.

#### Sex-linked genomic regions

Genomic regions with sex-linked loci were identified with RADsex (v.1.2, Feron et al. 2021). Briefly, demultiplexed raw reads of individuals with known sex were used to identify reads biased toward one sex. Analysis was performed for each species separately (*S. mentella*: N = 783 individuals; *S. fasciatus*: N = 479 individuals), with a minimum depth of five reads and a Bonferroni correction. Reads were then mapped to the *S. aleutianus* genome to identify genomic regions associated with sex. For *S. mentella*, 128 loci were identified as biased toward females on chromosome 3 (scaffold 72_5), in two blocs: 52 loci between positions 30,232,885 – 32,799,890 and 76 loci between positions 38,250,232 – 42,389,126 (Fig. S1). In addition, two other loci were identified as biased toward females, one in chromosome 16 (scaffold 62_5: position 5,835,562) and one in an unplaced scaffold (scaffold 73_2: position 9,695). For *S. fasciatus*, 13 loci identified on chromosome 3 between positions 30,657,861 – 31,523,318 were identified as biased toward females (Fig. S1). This region corresponds to a subset of the one also identified in *S. mentella*. SNPs identified within the sex-linked genomic regions were removed from species specific SNPs panels.

### Characterizing the genomic population structure for *Sebastes* spp

Three different SNPs datasets were used to explore *Sebastes* spp. inter and intra-specific genomic structure (Table 2). First, the *Sebastes* spp. dataset including the three *Sebastes* species was used to investigate inter-specific structure. Population structure was assessed using both principal component analysis (PCA) and clustering analysis. PCA was performed using the *glPCA* function of adegenet R package (Jombart 2008) and clustering analysis was performed using Admixture (v.1.3.0, Alexander et al. 2009) with K = 3 number of clusters. Finally, SnapClust (Beugin et al. 2018) was used to classify individuals as *S. mentella*, *S. fasciatus* or *S. norvegicus,* and to identify introgressed individuals.

SnapClust was first run on the complete *Sebastes* spp. dataset with K = 3 to classify individuals according to one of the three clusters (representing the three species). Then, the analysis was rerun on a subset dataset excluding the few *S. norvegicus* samples, with K = 2, and allowing for hybrid detections between *S. mentella* and *S. fasciatus*. Individuals identified as introgressed were further explored using the triangulaR package (Wiens et al. 2025) to assess how observed hybrid index and interclass heterozygosity matched theoretical expectations.

Species classification obtained using the *Sebastes* spp. dataset was used to generate species-specific datasets for *S. mentella* and *S. fasciatus* and explore intraspecific genomic structure (Table 2). PCAs were performed as described above for the species-specific datasets. Both Admixture and SnapClust were used to determine the number of genetic groups within each species. The optimal number of clusters (K) was obtained with cross-validation for Admixture, and with the *choose.cluster* function for SnapClust, testing K = 1 to 10 for each species. Indices of genetic differentiation (*F*_ST_) were computed among species, and among genetic groups within species using the dartR package (Gruber et al. 2018), with confidence intervals (CI) computed from 999 bootstraps.

### Spatiotemporal variation of genomic population structure

Temporal variation in genomic structure was examined by comparing the genetic composition of the 2010s cohorts to that of older cohorts within recent samples (2018-2021) and by characterizing the genetic composition of the 2000s cohort based on samples collected in 2005. Redfish from the 2010s cohorts were identified based on their fork length compared to known distribution in Unit 1 from stock assessments (2018: 20-23 cm, 2020: 22-24 cm, 2021: 23-25 cm; Senay et al. 2021, 2023). We also compared genetic composition of smaller fish (<28 cm) to larger ones in recent samples as a complementary approach to detect temporal patterns without relying strictly on cohort assignment. We used Chi-squared tests to assess whether the proportions of genetic groups varied according to size category or management units.

Spatial variation in the relative composition (proportion) of species or genetic groups in the recent samples was evaluated with respect to two key factors employed in Redfish management, namely, depth and management units. We fitted baseline-category multinomial logit models (Agresti 2002) using the mclogit R package (Elff 2024) with response variables defined as the proportion of (1) *S. mentella* and *S. fasciatus,* (2) each S*. mentella* genetic group, or (3) each *S. fasciatus* genetic group in a sample.

We built models of increasing levels of complexity, incorporating the effect of depth and management units (Unit 1, 1&2 and 2), as well as their interaction. Models were compared using Akaike’s information criterion corrected for sample size (AICc).

## Results

### Genomic population structure in the management units

#### Sebastes species

The *Sebastes* spp. dataset (i.e., all genotyped specimens, N =2,248; Table 2) was used to identify species and characterize introgression among them. All except one redfish from the 2000s cohort (2005 samples) were *S. fasciatus* (N = 792/793), while 62% of the redfish from the 2010s cohorts and contemporary fish (2018-2021 samples) were *S. mentella* (N = 680/1090; Fig. 2, Table 1). A total of 31 introgressed individuals between *S. mentella* and *S. fasciatus* were detected with SnapClust, including 3 F1 hybrids (i.e., 50% each species), 12 *S. mentella* backcrosses (i.e., 75% *S. mentella*) and 16 *S. fasciatus* backcrosses (i.e., 75% *S. fasciatus*), corresponding to individuals with intermediated values in PCA and Admixture plots (Fig. 2AB). Further evaluation using triangulaR suggested that while F1 hybrids were correctly identified, only two *S. mentella* and one *S. fasciatus* backcrosses matched the theoretical expected values of hybrid index and interclass heterozygosity (Fig. S2). The other identified backcrosses were characterized by higher rates of missing values or within the continuum of introgression between the two species. The *F*_ST_ reached 0.424 between *S. mentella* and *S. fasciatus* (*p* < 0.001, 95% CI [0.419, 0.428]), 0.234 between *S. mentella* and *S. norvegicus* (*p* < 0.001, 95% CI [0.229, 0.239]), and 0.505 between *S. fasciatus* and *S. norvegicus* (*p* < 0.001, 95% CI [0.500, 0.511]).

**Figure 2.**
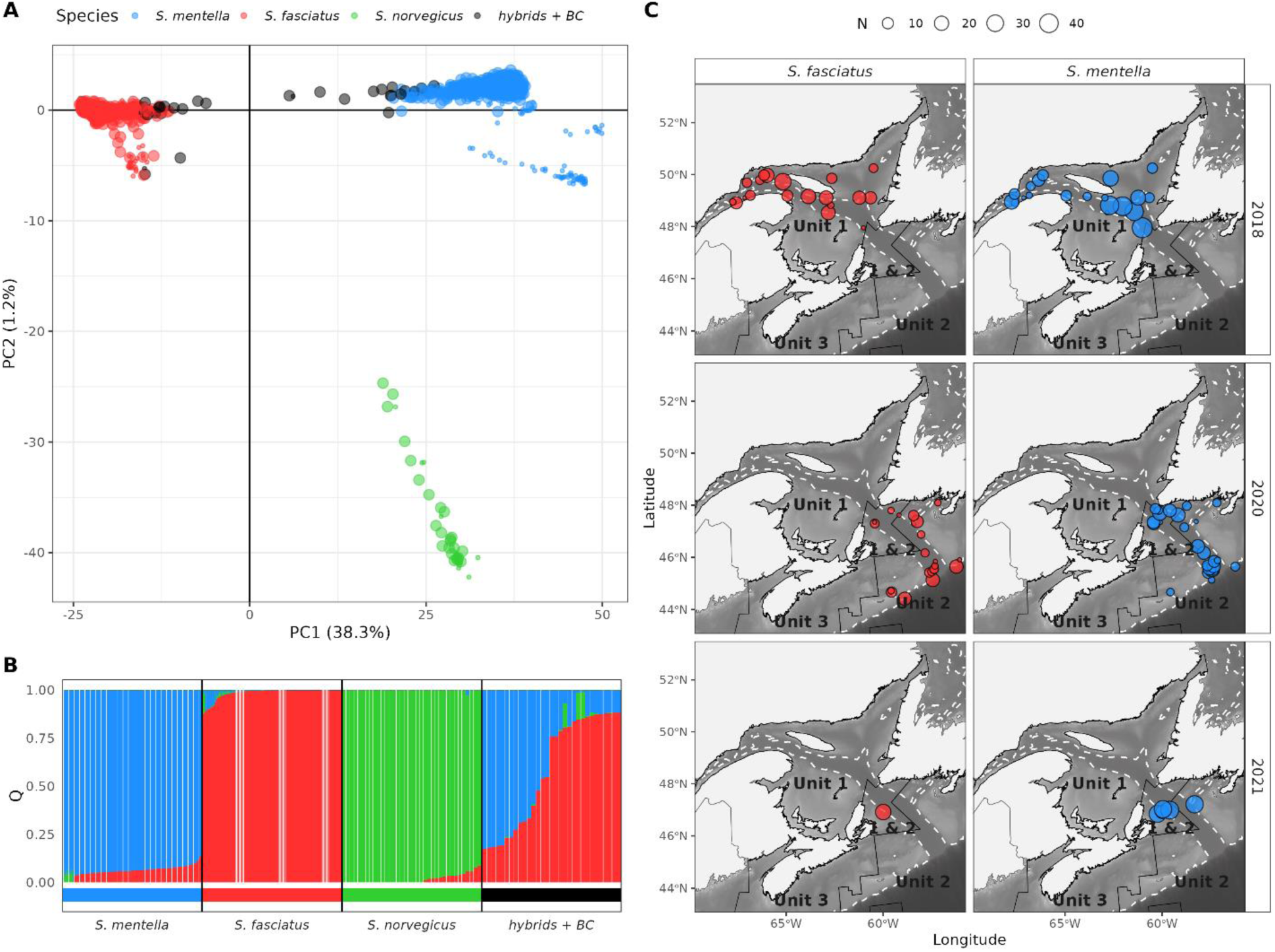
:Population structure using the *Sebastes* spp. dataset (N = 2,248), and distribution of *S. mentella* and *S. fasciatus* in the management Units 1, 1&2 and 2 from the recent samples (2018-2021, N = 1,066). (A) Principal component analysis of genetic variation observed at the individual level. The first two axes are presented, and larger dots denoted recent samples. (B) Membership probability (Q) observed at K = 3 for each species, and for individuals identified as introgressed. (C) Distribution of recent samples across years and management units. White dashed line represents the bathymetry at 300 m depth and point size reflected sample size. Color referred to the genetic groups (i.e., species) assigned by SnapClust at K = 3.

#### *Sebastes mentella* population structure

The analysis of the *S. mentella* dataset suggested the presence of five genetic groups. The first two axes of the PCA revealed five distinct clusters (Fig. 3A), a number that matched K = 5 optimal number of populations identified by both SnapClust and Admixture (Fig. S4). However, Admixture results at K = 5 or K = 4 matched only partially those of the PCA (Fig. 3B, S5). Thus, we interpreted PCA and SnapClust results together, which were clearer and coherent, to present population structure of *S. mentella* (Fig. 3A).

**Figure 3.**
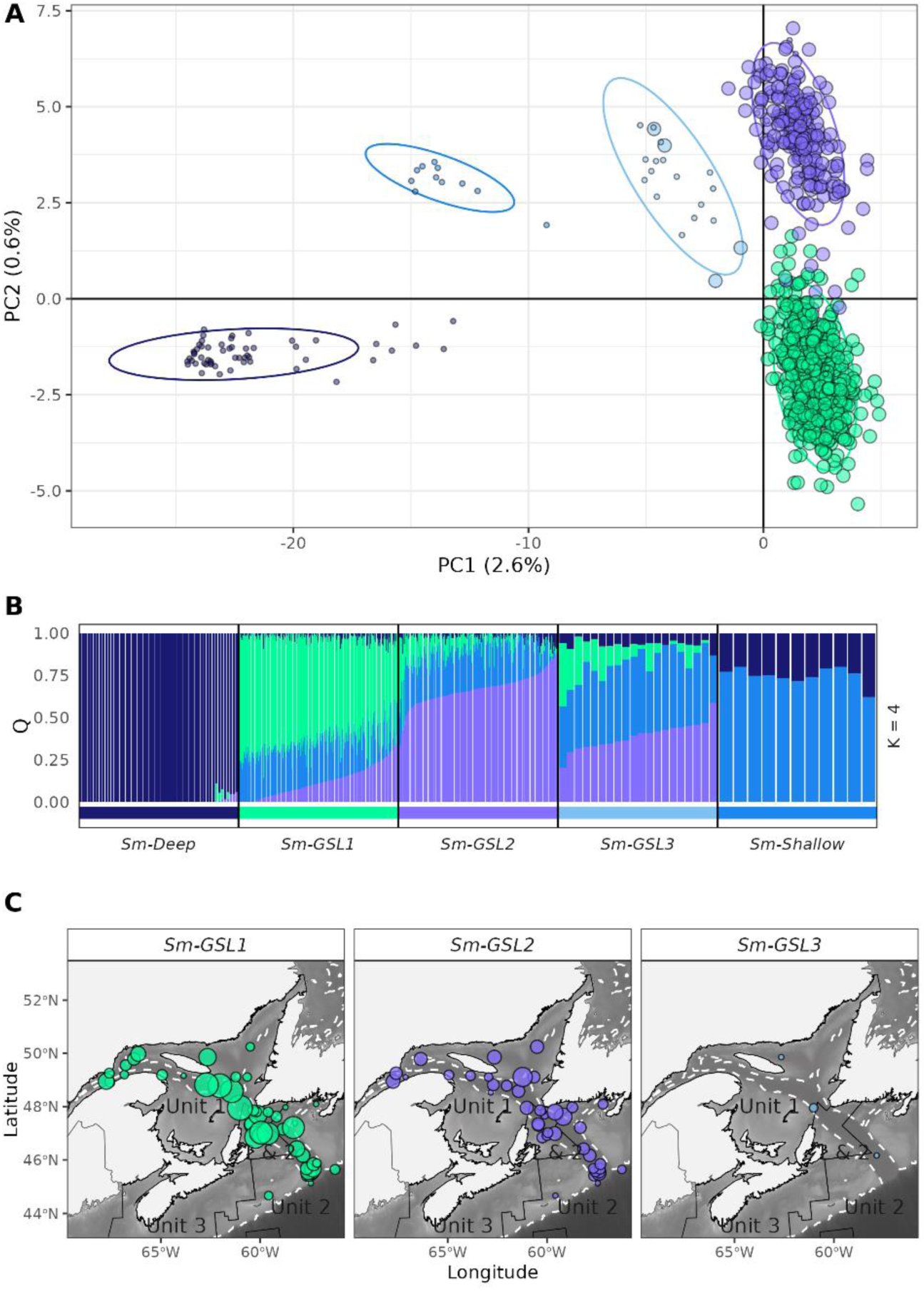
Population structure using the *S. mentella* dataset (N = 810), and distribution of genetic groups in the management Units 1, 1&2 and 2 in the recent samples (2018-2021, N = 705). (A) Principal component analysis of genetic variation observed at the individual level. The first two axes are presented, and larger dots denoted recent samples. B) Membership probability (Q) observed at K = 4 (as no other significant groups at the optimal K = 5, see Fig. S5) (C) Distribution of the genetic groups in recent samples across management units. White dashed line represents the bathymetry at 300 m depth and point size reflected sample size. Color referred to the genetic groups assigned by SnapClust at K = 5.

Two of the genetic groups observed correspond to the previously described ecotypes *S. mentella* “*Deep*” and “*Shallow*” (hereafter *Sm-Deep* and *Sm-Shallow*, respectively) and originated from samples collected outside the GSL-LC (References; Table 1, Fig. 1; Benestan et al. 2021). The third ecotype previously described as “*GSL*” was subdivided into three distinct groups which are henceforth referred to as *Sm- GSL1*, *Sm-GSL2*, and *Sm-GSL3*. Both *Sm-GSL1* and *Sm-GSL-2* represented the vast majority of *S. mentella* in Units 1, 1&2 and 2 in recent samples (N = 701/705; 2018-2021; Fig. 3C). The *Sm-GSL3* fish were rare (N = 4/705) in recent samples collected from the GSL-LC (Fig. 3C) but were more abundant in samples taken off southwestern Labrador in NAFO Division 2J (Fig. 1).

Genetic differentiation between *S. mentella* ecotypes was greatest between the *Sm-Deep* and the *Sm- GSL* ecotypes (considering the three genetic groups collectively) compared to other pairwise comparisons. The *F*_ST_ values were 0.094 between *Sm-Deep* and *Sm-GSL* (*p* < 0.001, 95% CI [0.092, 0.096]), 0.074 between *Sm-Deep* and *Sm-Shallow* (*p* < 0.001, 95% CI [0.071, 0.077]), and 0.061 between *Sm-Shallow* and *Sm-GSL* (*p* < 0.001, 95% CI [0.059, 0.063]). Within the *GSL* ecotype, the highest *F*_ST_ values (≥0.018) were reached with the *Sm-GSL3* and the two other *Sm-GSL* genetic groups (*Sm-GSL1*, 0.020, *p* < 0.001, 95% CI [0.019 0.021]; *Sm-GSL2*, 0.018, *p* < 0.001, 95% CI [0.017 0.019]). In contrast, the *F*_ST_ between *Sm-GSL1* and *Sm-GSL2* was 0.006 (*p* < 0.001, 95% CI [0.005, 0.006]). Based on the *Sebastes* spp dataset Admixture result, *Sm-Deep* exhibited on average 5.1 ± 1.0 % introgression with *S. norvegicus* while all three *S. mentella GSL* genetic groups presented on average 6.5 ± 2.0 % introgression with *S. fasciatus* (Fig. S6).

#### *Sebastes fasciatus* population structure

At least seven genetic groups were detected in the *S. fasciatus* dataset. The first five axes of the PCA revealed significant structure within *S. fasciatus* (Fig. 4A, Fig. S7A). The optimal number of genetic groups was K = 7 with SnapClust and K = 5 with Admixture (Fig. S7BC). A closer examination of Admixture membership probability of K = 5 and 7, coupled with SnapClust results, supported the presence of seven genetic groups (Fig. 4B). Two genetic groups were present mainly in reference samples, i.e., (1) *Sf-2J* was observed in NAFO Division 2J, except for five samples which were collected in NAFO Division 3P in recent samples, and (2) *Sf-BON* was only observed in Bonne Bay, a semi-closed area within NAFO Division 4R. Five genetic groups were observed in the 2010s cohorts and contemporary fish (2018- 2021): (1) *Sf-GSL1,* which is mainly observed in the GSL; (2) *Sf-GSL2,* more predominant in the GSL and the Cabot Strait; (3) *Sf-GSL3,* observed across the GSL-LC; (4) *Sf-FAN1*, mainly observed in the Laurentian Fan, along the continental margin; and (5) *Sf-FAN2,* mainly observed in the Laurentian Channel between the Cabot Strait and Laurentian Fan, and in the Gulf of Maine (Fig. 4C). Most of the fish collected in 2005, representative of the 2000s cohort (N = 787/792), captured in the GSL, belong to the Sf-FAN1 genetic group (Fig. S3). Note that due to their disproportionate representation and potential bias on structure analysis, these last individuals were excluded from the *S. fasciatus* dataset and analyzed separately.

**Figure 4.**
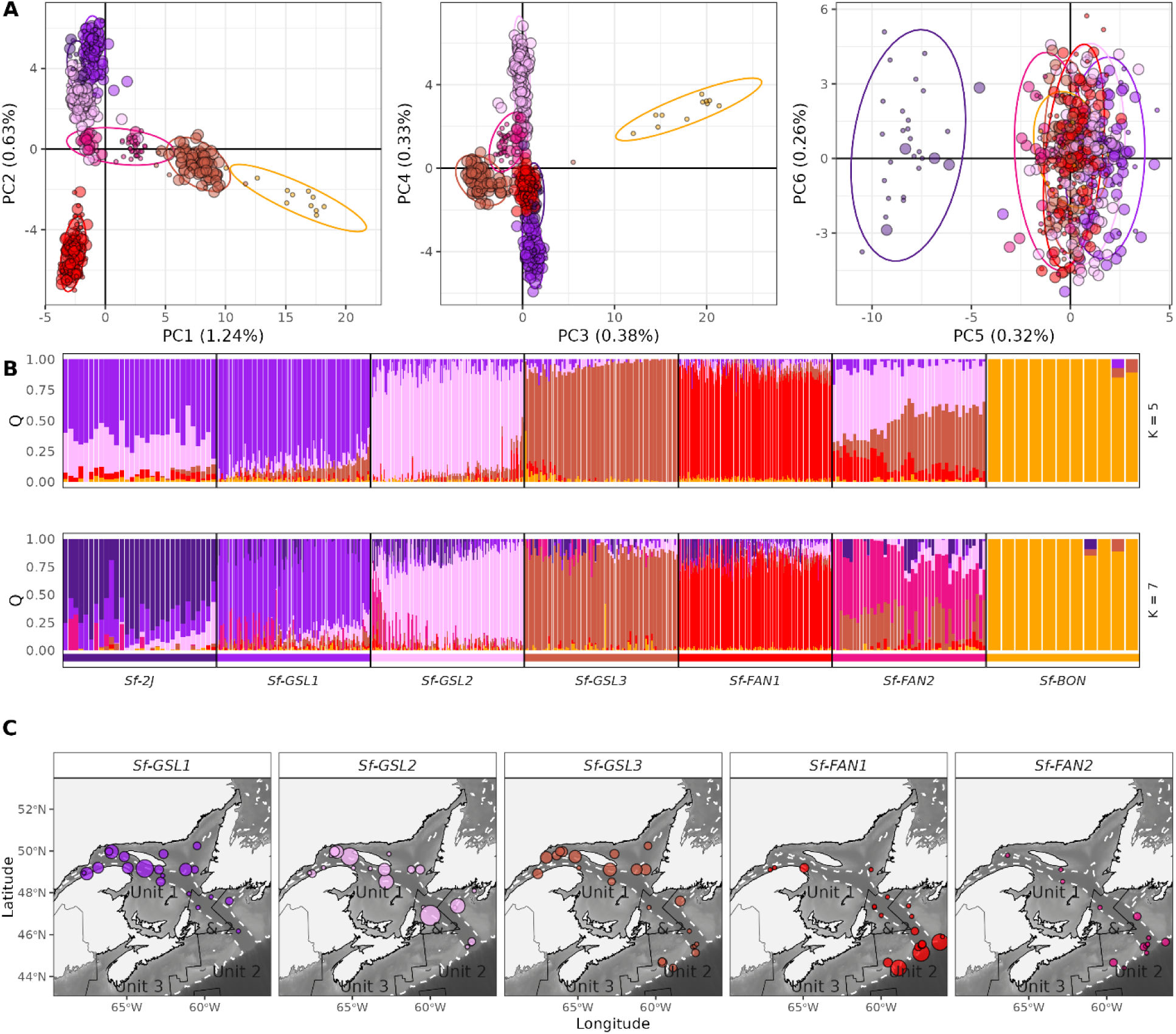
: Population structure using the *S. fasciatus* dataset (N = 561) and distribution of the main genetic groups in the management Units 1, 1&2 and 2 in the recent samples (2018-2021, N = 335). (A) Principal component analysis of genetic variation observed at the individual level. The first six axes are presented, and larger dots denoted recent samples. (B) Membership probability (Q) observed at Admixture K = 5 and 7. (C) Distribution of the main genetic groups in the recent samples. White dashed line represents the bathymetry at 300 m depth and sampling location size reflected sample size. Color referred to the genetic groups assigned by SnapClust at K = 7.

Highest *F*_ST_ values were observed with the *Sf-BON* cluster (all *F*_ST_ > 0.109, all p-values < 0.001). Otherwise, *F*_ST_ values among other genetic groups ranged from 0.009 (*Sf-GSL1* and *Sf-FAN2*) to 0.039 (*Sf- 2J* and *Sf-GSL3*; all p-values < 0.001). Two *S. fasciatus* genetic groups showed introgression with other *Sebastes* spp. based on membership probability (Q) provided by Admixture analysis. The genetic groups *Sf-GSL1* and *Sf-2J* showed introgression averaging 8.8 ± 1.4 % with *S. mentella* and of 11.0 ± 2.3 % with *S. norvegicus*, respectively (Fig. S8).

### Spatiotemporal genomic composition

We examined the fork length distribution (i.e., age proxy) for the 2010s cohorts and contemporary samples (2018-2021) to try to identify spatiotemporal variation in genomic structure (Fig. 5). Fish from the 2010s cohorts were identified based on length using expected lengths for these cohorts in each year from Unit 1 redfish stock assessments (Fig. 5A, grey bars). Almost half of the *S. mentella* (N =325, 46.4%) and a third of the *S. fasciatus* (N = 99, 29.4%) were attributed to the 2010s cohorts, and the rest were mainly larger fish. For *S. mentella*, 2010s cohorts were captured across the three sampling years and management units (Fig. 5A). For *S. fasciatus*, 2010s cohorts were mainly captured in Unit 2 in our sampling design, representing 48.2% of *S. fasciatus* in Unit 2 compared to 16.8% in Unit 1 (Fig. 5A).

**Figure 5.**
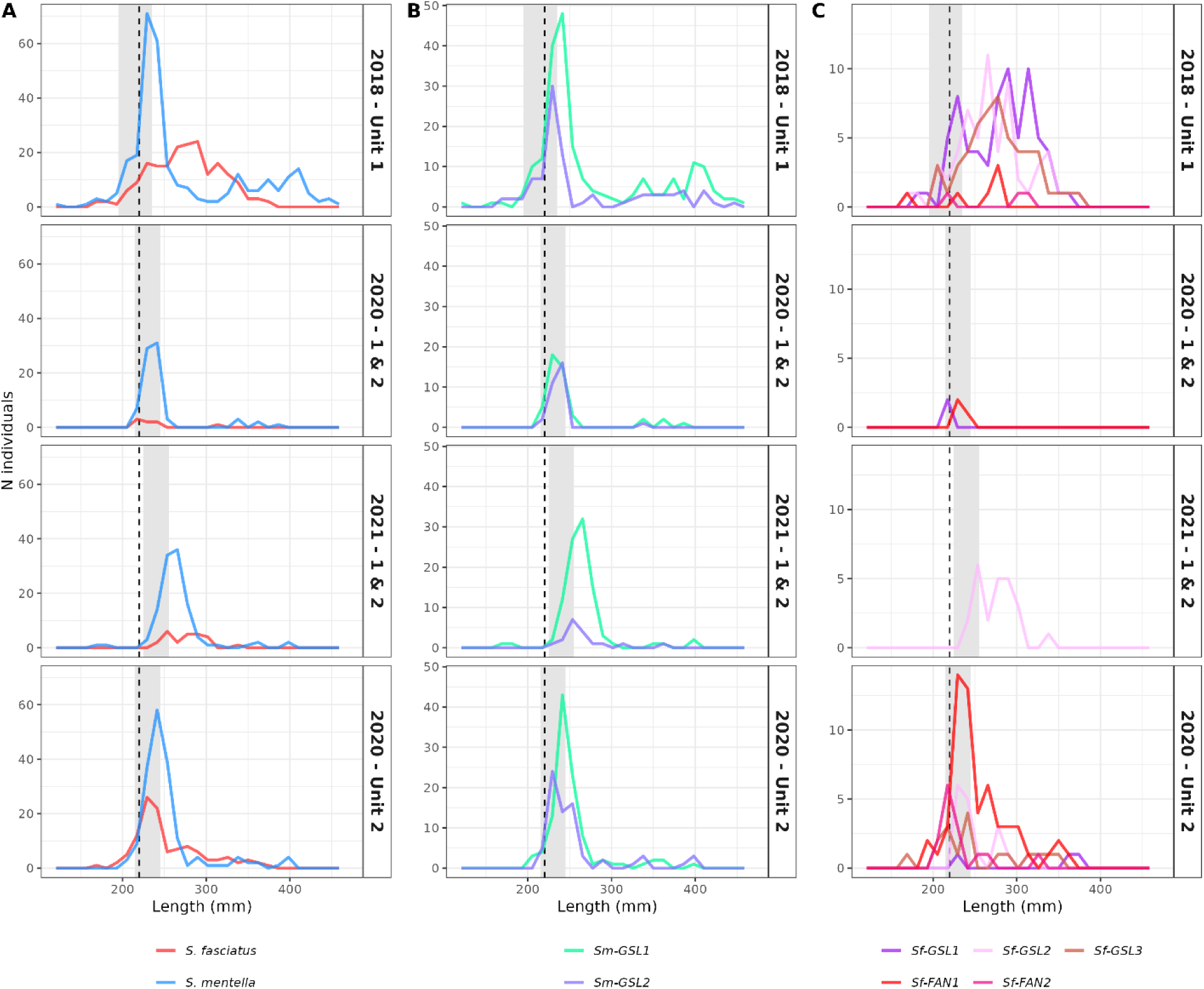
: Fork length distribution of *S. mentella* and *S. fasciatus* species and genetic groups across sampling years and management Units (1, 1&2, 2) in the recent samples (2018-2021). Panels present distributions (A) between species, (B) between *Sm-GSL1* and *Sm-GSL2* within *S. mentella*, and (C) between main genetic groups within *S. fasciatus*. Gray zones represent the expected body length for the 2010s cohorts based on stock assessments (Senay et al. 2021, 2023), while the dashed lines represent is the minimum length (220 mm) of sampled fish. Color referred to the genetic groups defined by SnapClust. Note that Unit 2 samples in 2021 are not included as only composed of a few *S. mentella* (N = 33).

We then examined fork length distributions by genetic group for each species within and across management units. For *S. mentella*, a similar length distribution was observed for both the *Sm-GSL1* and *Sm-GSL2* genetic groups within and across all management units (Fig. 5B). In contrast, *S. fasciatus* genetic groups showed distinct length distribution patterns between Units 1, 1&2 and 2 (Fig. 5C). Unlike *S. mentella*, no prominent peak associated with the 2010s cohorts was observed for the three main *S. fasciatus* genetic groups of Unit 1 (i.e., *Sf-GSL1*, *Sf-GSL2* and *Sf-GSL3*). Instead, these groups displayed multiple small peaks of different lengths (Fig. 5C). In Unit 2, the main recruitment of the 2010s cohorts was attributed to *Sf-FAN1*, while smaller peaks were noted for *Sf-GSL2*, *Sf-GSL3* and *Sf-FAN2*; Fig. 5C). In Unit 1&2, a single genetic group of *S. fasciatus*, *Sf-GSL2*, was mainly captured and its length distribution suggested a recruitment event largely aligned with the 2010s cohorts (Fig. 5C).

Comparisons of frequency distributions of genetic groups between younger and older cohorts (<28 cm, ≥ 28 cm) and across management units indicated variability solely for *S. fasciatus* (Fig. 6). For *S. mentella*, both *Sm-GSL1* and *Sm-GSL2* genetic groups were present in similar proportions in smaller and larger fish (χ² = 2.49, df = 2, p = 0.29), and among units (χ² = 8.33, df = 4, p = 0.08), with some variation most likely due to sampling variability (Fig. 6A). For *S. fasciatus*, differences in genetic composition varied slightly between the two body length classes (χ² = 24.83, df = 5, p < 0.001; Fig. 6B). In contrast, genetic composition differed substantially among management units (χ² = 192.69, df = 10, p < 0.001; Fig. 6B). The Unit 1 was dominated by the three *Sf-GSL* groups, most notably *Sf-GSL1*. *S. fasciatus* from Unit 1&2 were relatively rare, but were primarily from the *Sf-GSL2* genetic group, a group present across all units. In Unit 2, a greater diversity of genetic groups was observed, although *Sf-FAN1* predominated among both smaller and larger fish.

**Figure 6.**
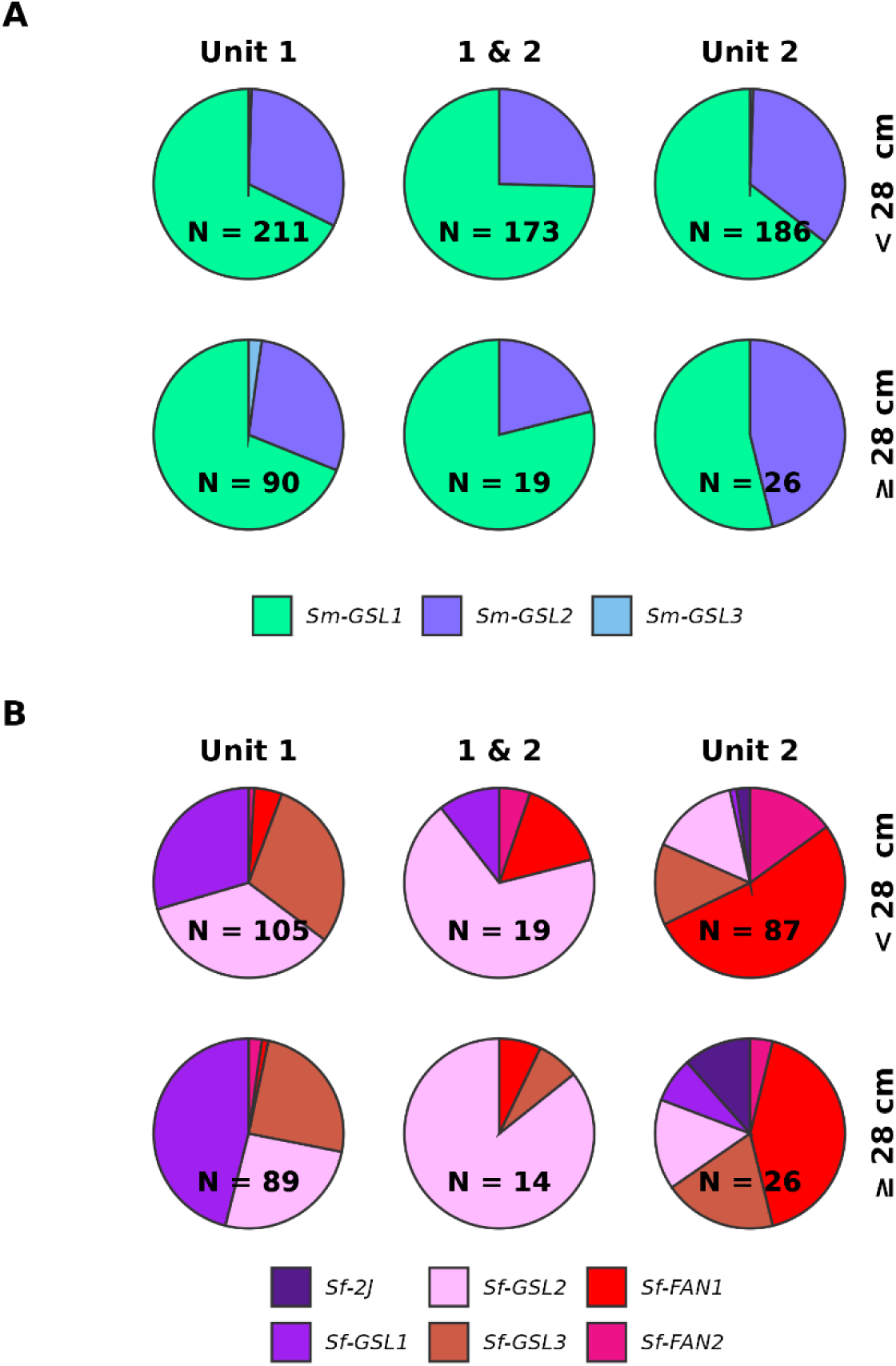
: Genetic group composition by management units (Unit 1, 1&2, 2) for smaller (fork length <28 cm) and larger (fork length ≥ 28 cm) fish of (A) *S. mentella* and (B) *S. fasciatus* in the recent samples (2018- 2021). Color referred to the genetic groups defined by SnapClust.

### Three-dimensional spatial structure

We used baseline-category multinomial logit models to explore how population structure relates to depth and management units from the 2010s cohorts and contemporary samples (Fig. 7). At the species level, the best model based on AICc included an interaction between depth and management units, with delta AICc ≥ 28.8 relative to the four other models (Table S1A). As expected, the predicted proportions of *S. mentella* in a sample increased with depth while the opposite trend was observed for *S. fasciatus* (Fig. 7A, Table S1B). The slope of the relation with depth was less steep in Unit 2, compared to Units 1 and 1&2.

**Figure 7.**
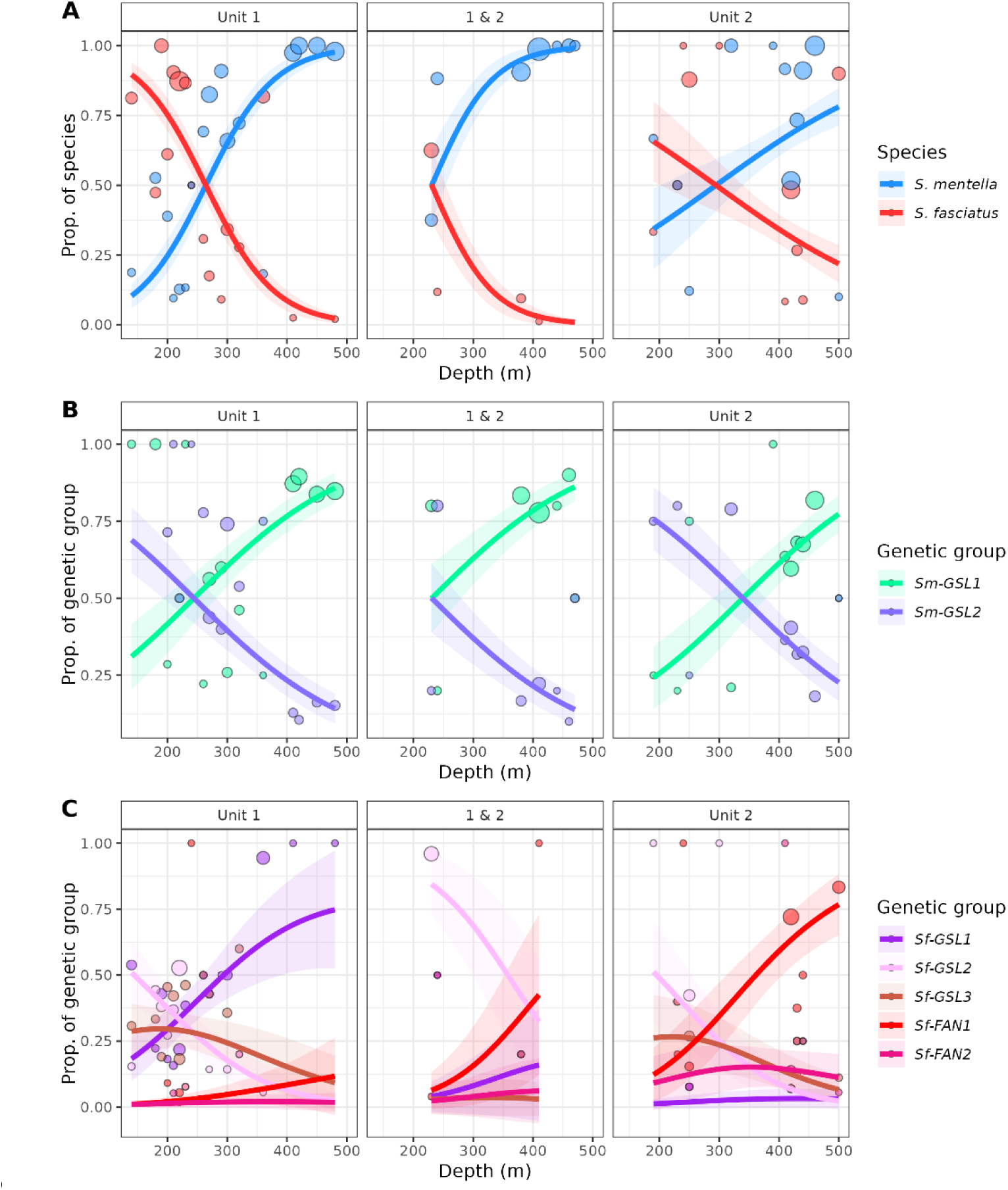
: Relative composition of (A) species (N = 1045), (B) *S. mentella* (N = 701) and (C) *S. fasciatus* (N = 335) genetic groups in recent samples (2018-2021), with respect to depth and management units (Unit 1, 1&2, 2). Predictions are derived from baseline-category multinomial logit models where best models were selected by AICc (Tables S1-3). Lines represented the best model predictions, shaded areas 95% CI, circles the observed proportions in bin depth of 10 m and color the genetic groups defined by SnapClust.

For the two main genetic groups within *S. mentella*, the best model based on AICc also included both depth and management units, with delta AICc ≥ 2.6 relative to the four other models (Table S2A). In this model, the proportion of *Sm-GSL1* increased with depth, while *Sm-GSL2* showed the opposite trend (Fig. 7B, Table S2B). Moreover, the proportion of GSL-1 was lower in Unit 2 compared to Units 1 and 1&2.

For the five main genetic groups within *S. fasciatus*, the best model based on AICc included both depth and management units, with delta AICc ≥ 5.4 relative to the four other models (Table S3A). Two genetic groups, *Sf-GSL1* and *Sf-FAN1*, were observed in higher proportion in deeper samples (Fig. 7C). These relationships with depth were only present in the management unit where these groups were more abundant, specifically in Unit 1 for *Sf-GSL1* and in Unit 2 for *Sf-FAN1* (Fig. 7C, Table S3B). Two other genetic groups, *Sf-GSL2* and *Sf-GSL3*, were observed in higher proportion in shallower samples (Fig. 7C). In these models, the proportion of *Sf-GSL2* and *Sf-GSL3* were, respectively, higher and lower in Unit 1&2 compared to Units 1 and 2 (Fig. 7C, Table S3B). Lastly, for the *Sf-FAN2* genetic group, the proportion predicted in each management unit was always low (Fig. 7C, Table S3B).

## Discussion

In this study, we characterized the population genomics of *S. mentella* and *S. fasciatus* at fine scale in the GSL-LC. We confirmed admixture between the two species but also identified admixture of the two species with *S. norvegicus*, and revealed more population structure for both species than previously reported due to the finer scale study. We also improved our understanding of the distribution of the numerous *S. fasciatus* genetic groups. Both species and their intraspecific genetic groups were distributed non-randomly across depth and management units. Our results also indicate difference in the length distribution across *Sebastes* genetic groups and management units, suggesting spatiotemporal variability of recruitment between species and their intraspecific groups. We will discuss these results and how the complex three-dimensional genomic structure in redfish for the GSL-LC underscored the need for management approaches that go beyond the current units and depth-based regulations to ensure the long-term sustainability of all *S. mentella* and *S. fasciatus* populations.

### Mechanisms driving genomic structure and diversification

#### Introgressive hybridization

The exchange of genetic material between species through hybridization and repeated backcrossing, known as introgressive hybridization, plays a significant role in shaping genetic diversity within marine environments. In the Northwest Atlantic, secondary contacts, the renewed genetic exchanges following isolation in marine refugia, has been documented in many species during post-glacial recolonization (Maggs et al. 2008), including between *S. fasciatus* and *S. mentella* (Roques et al. 2001; Benestan et al. 2021). Within the GSL-LC, the presence of introgressive hybridization appears to be one of the distinctive features of species and populations divergence. For S. *mentella*, all three genetic groups or populations within the *GSL* ecotype present an admixture signal with *S. fasciatus* (Fig. S6B), suggesting that introgressive hybridization occurred historically, prior the divergence of this ecotype into populations. In *S. fasciatus*, introgressive signals with *S. mentella* were detected only in the *Sf-GSL1* population (Fig. S8A). We also found evidence of introgression between *S. norvegicus*, the third redfish species encountered in the Northwest Atlantic, and both *S. mentella Deep* (Fig. S6A), and *S. fasciatus Sf- 2J* (Fig. S8B). Similar patterns of introgression between *S. mentella* and *S. norvegicus* have been previously reported in the Northeast Atlantic (Pampoulie and Daníelsdóttir 2008; Saha et al. 2017b).

Admixed *Sebastes* genetic groups are consistently found at the edges of their species distributions in our study. Specifically, the *Sm-GSL* ecotype is located at the southern limits of *S. mentella* range (Benestan et al. 2021; Cadigan et al. 2022). Conversely, the introgressed populations of *S. fasciatus* are observed both within the GSL (*Sf-GSL1*) and along the Labrador Coast (*Sf-2J*; see also Benestan et al. 2021), marking the northernmost extent of *S. fasciatus* range. Similarly, introgression is associated to species depth limits: the *Sm-Deep* ecotype inhabits deeper waters, aligning with the described habitat of *S. norvegicus*, while the *Sf-GSL1* population is observed at depths typical of *S. mentella* within the Gulf of St. Lawrence. Given the presumably rare and historical hybridization and introgression among *Sebastes*, we hypothesize that introgressed individuals may exhibit enhanced fitness at the margin of species distribution. Hybridization can play an important role in adaptation (Seehausen 2004; Hedrick 2013), potentially giving rise to novel or transgressive phenotypes capable of facilitating range expansion (Pfennig et al. 2016). What stands out in the Northwest Atlantic is the repeated occurrence of this process across the three *Sebastes* species, suggesting that introgression may promote diversification at range boundaries. Beneficial genetic variants acquired through introgressive hybridization therefore appear to represent an important evolutionary mechanism for adaptation in *Sebastes*.

Despite the strong historical signal of introgressive hybridization, we detected very few contemporary hybrids. Consistent with earlier work (e.g., Saha et al. 2017b; Benestan et al. 2021), first-generation hybrids and backcrossed individuals appear to be rare. In our study, only three putative hybrids were identified, representing 0.1% of all genotyped individuals. Although sympatric areas like the GSL-LC, combined with our extensive sampling, would have maximized the likelihood of detecting hybrids if they were present, the extremely low number observed indicates that contemporary gene flow between *S. mentella* and *S. fasciatus* is very limited. This is further supported by the pronounced genetic differentiation maintained between species. Nevertheless, the accurate identification of hybrids and backcrossed individuals remains critical, underscoring the importance of including all sympatric *Sebastes* species in genetic analyses.

#### Behavioral and ecological barriers

The high number of genetically distinct populations of *S. fasciatus* within the GSL-LC is noteworthy and contrasts with the lower population differentiation observed in *S. mentella* within the same areas. This pattern raises questions about the mechanisms that generate such diversity and maintain differentiation among *S. fasciatus* populations in a relatively small area. *Sebastes* are recognized as a prominent example of adaptive radiation in the Pacific, with nearly a hundred species described (Hyde and Vetter 2007). Therefore, observing high levels of diversification among redfish in the Atlantic, particularly given their relatively recent arrival (Shum et al. 2015), is consistent with the evolutionary capacity of the genus for rapid diversification. Difference between *S. mentella* and *S. fasciatus* in terms of structure complexity may be partly explained by differences in mobility and life history strategies between species. Limited adult movement tends to reinforce differentiation among groups. For instance, in Pacific *Sebastes*, populations exhibiting lower dispersal rates and higher site fidelity tend to display greater population structure (Andrews et al. 2018). Based on body shape analysis, *S. fasciatus* is less fusiform, a morphology associated with a more sedentary behavior compared to *S. mentella* (Valentin et al. 2006). In contrast, *S. mentella* exhibits relatively high mobility, with documented movements between nursery and adult grounds in the Barents Sea (Drevetnyak and Nedreaas 2009).

The depth segregation among redfish populations within the GSL-LC can act as a pre-zygotic barrier to gene flow, further promoting diversification. Depth-related isolation has played a central role in the diversification of *Sebastes* (Ingram 2011; Shum et al. 2014; Behrens et al. 2021). In the present study, the close association between specific genetic groups and particular depth ranges supports the central role of depth in promoting reproductive isolation within the GSL-LC. As depth cannot explain all the observed structure, additional factors such as mate choice or temporal reproductive isolation could also play a role (Helvey 1982; Buonaccorsi et al. 2011; Johansson et al. 2012; Yamaguchi et al. 2025).

### Three-dimensional spatial population structure in the GSL-LC

#### *S. mentella GSL* ecotype subdivided into three populations

Complexity within *S. mentella* has for a long time been the subject of research and debate (Cadrin et al. 2010). In the Northeast Atlantic, three ecotypes are currently defined (*Deep*, *Shallow* and *Slope*), and recent genomic studies suggested more structure within most ecotypes (Saha et al. 2021; Jansson et al. 2025). In the Northwest Atlantic, genetic distinctiveness of *S. mentella* from the GSL were recognized for a long time (Roques et al. 2001; Valentin et al. 2014; Benestan et al. 2021). The *Sm-GSL* ecotype has been reported within the GSL-LC as well as in the NAFO Divisions 2G and 3K, with the hypothesis that juvenile movements occur through the Strait of Belle Isle (Benestan et al. 2021). Our results, with more individuals genotyped, suggested substructure within the *GSL* ecotype, with three distinct populations. The population mostly abundant outside the GSL-LC in 2J, *Sm-GSL3*, exhibits the highest genetic differentiation from the two other *Sm-GSL* populations. Re-examination of previous studies revealed some hints of this subdivision of the *Sm-GSL* ecotype into more components (see Fig. S3 in Benestan et al. 2021) and of genetic variance among sampling sites (Valentin et al. 2014). The hierarchical nature of the *Sebastes* structure combined with limited power (i.e., smaller sample size and fewer markers) could explain why this substructure could only be identified in our study.

Intraspecific differences in biological characteristics and life-history traits of *S. mentella* in the GSL-LC have been documented by several studies. While body shape appeared consistent across diverse sampling sites (Valentin et al. 2014), a parasitological study observed differences in infectious status between Unit 1 and Unit 2, which was interpreted as a possible indicator of population structure (Marcogliese et al. 2003). More recently, an analysis of otolith elemental fingerprint of *S. mentella* from the 2010s cohorts suggested the presence of two chemically distinct natal sources, following an east-west gradient across the northern GSL (Coussau et al. 2023). These chemically defined groups may correspond to the *Sm-GSL1* and *Sm-GSL2* populations observed in our study. Here we found that the proportion of each population varied with depth, an environmental component that could be related to differences detected via otolith chemistry. Furthermore, the proportions of *Sm-GSL1* and *Sm-GSL2* also showed slight variation across management units, which are themselves organized along an east-west axis. Further analyses integrating genetic, ecological, and chemical data will be necessary to clarify how these distinct lines of evidence relate to one another and contribute to our understanding of *S. mentella* population structure in the GSL-LC.

#### At least five *S. fasciatus* populations

Our findings reveal a complex pattern of population structure for *S. fasciatus* within the GSL, with at least five distinct genetic groups that exhibit a non-random three-dimensional distribution across the region. Previous genetic and morphometric studies identified variance and structure in this region (Valentin et al. 2014; Benestan et al. 2021). For instance, Benestan et al. (2021) reported at least two populations based on four sampling sites within the GSL-LC. Here, with a denser sampling scheme and greater sample size, our study identified five populations displaying distinct distributions between management units: three were abundant in Unit 1, while four were observed in Unit 2. Notably, the two populations associated with deeper environments, *Sf-GSL1* and *Sf-FAN1*, were each almost exclusively found within a single management unit. While a deeper *S. fasciatus* had previously been described in the Laurentian Fan (i.e., the edge of the Laurentian Channel; Valentin et al. 2014), this study is the first to characterize a comparable deepwater population within the GSL itself.

The two other predominant populations in Unit 1, *Sf-GSL2* and *Sf-GSL3*, were associated with shallower environments. Only one population, *Sf-FAN2*, did not display any clear spatial or depth association. This group included relatively few individuals, mainly in Units 1&2 and 2, and was genetically similar to “Reference” individuals from the Gulf of Maine. However these individuals were only clustered with the Gulf of Maine individuals according to Snapclust. In contrast, the PCA and Admixture analyses point to the presence of two sub-groups. These results suggest a more complex genetic structure, potentially reflecting the existence of an additional, uncharacterized population of *S. fasciatus* between the Laurentian Fan and the Gulf of Maine (Fig. 1). Some shallow and coastal areas were undersampled in our study, potentially hiding further *S. fasciatus* populations beyond those identified in this area. Increasing sample size and coverage, especially in these zones, could improve the resolution and characterization of biological units for this species. Nonetheless, current data, together with previous work (Benestan et al. 2021), support the likelihood of more populations than currently described.

#### Temporal variability of genomic composition

We assessed temporal variability by comparing samples collected at different times and by analyzing cohorts using fork length as a proxy. Using both approaches, we evaluated the contribution of populations to redfish recruitment and survival, and examined how genomic population structure varies over time in relation to episodic recruitment events. Our findings corroborate previous observations showing that GSL redfish from the 2010s cohorts were predominantly *S. mentella* and from the 2000s cohort consisted almost exclusively of *S. fasciatus* (Valentin et al. 2015; Brassard et al. 2017).

Beyond this difference in species composition between successive recruitment events, our findings also reveal variability in population contribution to these events but only for *S. fasciatus*. For *S. mentella*, both populations contributed equally to the current large biomass in the GSL-LC, suggesting that they both benefited from similarly favorable environmental conditions during the strong episodic recruitment (Burns et al. 2021). In contrast, our results for *S. fasciatus* with specimens from different cohorts indicate unequal contributions of populations to recent biomass increases across the study area. In Unit 1, the proportion of the 2010s cohorts fish in recent samples was smaller than that of older fish compared to Units 1&2 and 2 (Fig. 5A,C), suggesting variation in the strength of recruitment events across populations and management units. We also observed that all *S. fasciatus* collected in Unit 1 in 2005, and representing the 2000s cohort, exhibited a genetic composition almost exclusively corresponding to *Sf-FAN1*, a striking contrast to the genetic composition of *S. fasciatus* in Unit 1 in the 2010s cohorts and contemporary samples, dominated by *Sf-GSL1*, *Sf-GSL2*, and *Sf-GSL3* genetic groups. This result is consistent with that from Valentin et al. (2015), where, using microsatellite markers and a limited number of samples, the 2000s cohort had a genetic signature similar to *S. fasciatus* found outside the GSL-LC. Both approaches show spatiotemporal variability in *S. fasciatus* genomic composition.

Three hypotheses may explain the temporal variability of Sf-FAN1 population observed in Unit 1. The first, proposed by Valentin et al. (2015), suggests that individuals from external populations disperse to the GSL to use it as a spawning and nursery area. Alternatively, larvae originating from the Laurentian Fan may drift into the GSL, which would similarly function as a nursery area. For both hypotheses, the subsequent disappearance of large *S. fasciatus* cohorts after ca. 5-7 years in Unit1 may be either attributed to a migration outside of the GSL or the low survival in local conditions. A third hypothesis posits high recruitment of the *Sf-FAN1* population within the GSL individuals, followed by population-specific mortality between juvenile and adult stages. All hypotheses remain plausible, our results do not strongly support any of the three. A key limitation of our sampling is the absence of a well design temporal monitoring of genomic composition. Genotyping the earlier life stages of the 2010s cohort (i.e., at a younger age, before 2018) might have revealed a different genomic composition in Unit 1, e.g., a larger proportion of the *Sf-FAN1* population. To clarify these dynamics, longitudinal studies tracking successive cohorts across all life stages, including larvae, would be valuable, particularly when combined with approaches such as otolith microchemistry to assess the origin of captured individuals (Coussau et al. 2023). Such studies could elucidate movement patterns, recruitment sources, and population-specific contributions to biomass, thus providing a more comprehensive picture of population dynamics, connectivity and temporal variability in genetic composition for redfish in the GSL-LC.

We used fork length distributions as a proxy of age but this metric is influenced by several biological and technical factors inherent to our sampling design. For example, in Unit 1, the 2011–2013 cohorts exhibited slower growth rates (Senay et al. 2023) and earlier onset of reproduction (Brûlé et al. 2024) compared to the 1980s cohorts, assumed as larger fish in our dataset. Thus, fork length provides a reliable distinction between older (1980s) and younger (2010s) cohorts, as their size ranges do not overlap due to the reduced growth of the younger fish. Fork length distribution may also be affected by differences in fishing gear. Our results show a shift in the distribution of both species between 2020 and 2021 in Unit 1&2. This change likely reflects fish growth but also gear selectivity associated with distinct fishing events. Commercial fishery samples in Unit 1&2 tend to include larger fish (2021) than those from the scientific survey (2020), which may be partially explained by differences in gear selectivity and fishermen’s behavior (Fig. 5ABC). Despite these confounding factors, using fork length to assess genomic population contributions to recruitment remains the best approach given the available samples, although we acknowledge potential biases.

### Conclusion and science advices for an improved management of a complex population structure for *Sebastes*

The recent reopening of the redfish fishery in 2024 in Unit 1, following nearly three decades of moratorium, underscores the urgent need for management strategies that reflect the underlying biological complexity of *Sebastes* populations. Management measures tied to depth-related regulations were suggested to account for the relative abundance of *S. mentella* and *S. fasciatus* (DFO 2024), a pragmatic approach given morphological similarities between species. However, our results indicate that management units and depth alone do not fully capture this complexity. For instance, multiple populations of *S. fasciatus* occur across the management units. In addition, the distinct depth distribution of *S. fasciatus* populations would increase fishing pressure on those observed in deeper habitats. Our fine-scale genomics results suggest the current division of the GSL-LC into two management units does not correspond to the scale of population structure for both species. Ideally, management units would be delineated to reflect biological units but achieving a sensible alignment between both unit types remains a challenge in practice (Cadrin 2020). Maintaining genomic diversity within species is essential for their resilience and long-term sustainability. For *S. mentella*, the species exhibits broad-scale genomic structure over the study area, so combining Units 1 and 2 as a single management unit could be considered a reasonable approach, as previously advocated (Valentin et al. 2014). In contrast, the finer-scale genomic structure of *S. fasciatus* suggests that further subdivisions of management areas may be necessary to effectively maintain the species genetic diversity. This situation where population genomics results for multiple species suggest distinct management unit scales is not unique to the GSL-LC *Sebastes* species. In the Pacific, five *Sebastes* species of the Puget Sound areas showed contrasting population structure, and thus species-specific scales of management were recently recommended (Wray et al. 2025). Similarly, our study reveals that the two redfish species in the GSL-LC exhibit markedly different patterns and scales of population structure, emphasizing that a one-size-fits-all management strategy may be inadequate. Acknowledging all the redfish populations within the GSL- LC and adapting management measures to account for their spatial and depth distribution would be necessary. For instance, determining whether NAFO Subdivisions best capture the *S. fasciatus* population structure should be addressed in a future study.

It is also important not to assume that all populations of a *Sebastes* spp. contribute equally during massive recruitment events within the GSL-LC. Our results indicate uneven contributions among populations, particularly for *S. fasciatus*. Therefore, we recommend more rigorous monitoring of cohort genomic composition throughout life history, including through larval sampling. Such efforts will enable a clearer understanding of juvenile recruitment in relation to the composition of the spawning stock, and ultimately provide deeper insights into the dynamics of the different populations. To enable efficient genetic monitoring, we recommend developing a GT-seq SNP panel tailored to these populations. A similar panel has already been validated for several Pacific *Sebastes* species and perform reliably even among closely related taxa (Baetscher et al. 2023), with levels of differentiation comparable to those observed among *S. mentella* and *S. fasciatus* populations.

In conclusion, recognizing and monitoring the fine-scale population structure in *Sebastes* species is essential to their sustainable management in the GSL-LC and elsewhere. Our study adds to a growing body of literature demonstrating that cryptic population structure is a common, and often overlooked, feature of *Sebastes* fisheries worldwide (Berntson and Moran 2009; Saha et al. 2017a; Longo et al. 2022; Wray et al. 2025). By integrating a comprehensive picture of *Sebastes* population structure, including both inter- and intraspecific genetic diversity, into fisheries management, we can more effectively safeguard biodiversity and ensure the long-term productivity of these valuable resources. Similar approaches have proven essential for ensuring the sustainability of economically valuable species (e.g., Atlantic salmon *Salmo salar*, (Lehnert et al. 2023) and a similar rational should be applied to all ecologically important species.

## Acknowledgment

We thanks the Atlantic Groundfish Council for sampling of Unit 2 and the devoted personal from the NGSL survey for years of collaboration. Laboratory work would not have been possible without the contribution of Gregoire Cortial, Gabriel Bardaxoglou and Jade Larivière. We acknowledge the use of AI-based language models (i.e., GPT-5 and GPT-5 Nano) to assist with correction of portions of the manuscript and paraphrasing, aiming to improve its clarity and accuracy in English. All content generated or edited by these models was thoroughly reviewed and verified, and we take full responsibility for its integrity and accuracy. We dedicate this work to the memory of Eric Parent, who made invaluable contributions to our understanding of redfish structure over the past decades, and who left us just a short time before his retirement. Throughout his career, he contributed to the advancement of population genetics, from its early reliance on allozymes to the adoption of microsatellites and, more recently, SNP markers. His contributions to our understanding of the diverse marine species of the Gulf of St. Lawrence are immeasurable, and his insight and presence are deeply missed.

## Data availability statement

Raw sequence data for the ddRAD datasets will be available in the Sequence Read Archive (SRA) under the BioProject accession no. PRJNA1208513. The scripts to obtain SNP panels from raw reads and perform analyses will be available on Zenodo.

