## Supplementary Material for "Population genomics reveals fine-scale three-dimensional structure within two sympatric *Sebastes* species in the Northwest Atlantic"

**Table S1 (A) Comparison of candidate models by corrected Akaike Information Criteria (AICc) and (B) best model estimates for the change in proportions of *S. mentella* and *S. faciastus* across depth and management units (Units 1, 1&2, 2) in recent samples (2018-2021, N = 1045).**

| (A) Model | AICc | dAIC | df |
|---|---:|---:|---:|
| **Depth * Unit** | **997.9** | **0.0** | **6** |
| Depth + Unit | 1026.7 | 28.8 | 4 |
| Depth | 1092.0 | 94.1 | 2 |
| Unit | 1276.4 | 278.6 | 3 |
| Null | 1320.5 | 322.6 | 1 |

| (B) Variable | Estimate | s.e. | z-value | p-value |
|---|---:|---:|---:|---:|
| Intercept | -4.516 | 0.888 | -5.09 | < 0.001 |
| Depth | 0.020 | 0.003 | 6.37 | < 0.001 |
| Unit 1 | -0.076 | 0.996 | -0.08 | 0.939 |
| Unit 2 | 2.690 | 1.070 | 2.51 | 0.012 |
| Depth x Unit 1 | -0.002 | 0.003 | -0.63 | 0.527 |
| Depth x Unit 2 | -0.013 | 0.003 | -3.92 | < 0.001 |

**Table S2 (A) Comparison of candidate models by corrected Akaike Information Criteria (AICc) and (B) best model estimates for the change in proportions of *S. mentella GSL1* and *GSL2* genetic groups across depth and management units (Units 1, 1&2, 2) in recent samples (2018-2021, N = 701).**

| (A) Model | AICc | dAICc | df |
|---|---:|---:|---:|
| **Depth + Unit** | **818.1** | **0.0** | **4** |
| Depth * Unit | 820.8 | 2.6 | 6 |
| Depth | 830.4 | 12.2 | 2 |
| Unit | 868.9 | 50.7 | 3 |
| Null | 871.1 | 52.9 | 1 |

| (B) Variable | Estimate | s.e. | z-value | p-value |
|---|---:|---:|---:|---:|
| Intercept | 1.761 | 0.436 | 4.04 | < 0.001 |
| Depth | -0.008 | 0.001 | -7.02 | < 0.001 |
| Unit 1 | 0.104 | 0.219 | -0.48 | 0.634 |
| Unit 2 | 0.831 | 0.231 | -3.59 | < 0.001 |

**Table S3 (A) Comparison of candidate models by corrected Akaike Information Criteria (AICc) and (B) best model estimates for the change in proportions of *S. fasciatus* main genetic groups across depth and management units (Units 1, 1&2, 2) in recent samples (2018-2021, N = 335).**

| (A) Model | AICc | dAICc | df |
|---|---|---|---|
| **Depth + Unit** | **800.9** | **0.0** | **16** |
| Depth * Unit | 806.4 | 5.4 | 24 |
| Depth | 846.6 | 45.1 | 12 |
| Unit | 886.6 | 85.6 | 8 |
| Null | 1018.4 | 217.4 | 4 |

| (B) Genetic group proportion | Variable | Estimate | s.e. | z-value | p-value |
|---|---|---|---|---|---|
| ***Sf-GSL2* vs *Sf-GSL1*** | Intercept | 6.054 | 1.111 | 5.45 | < 0.001 |
| | Depth | -0.013 | 0.003 | -4.60 | < 0.001 |
| | Unit 1 | -3.204 | 0.794 | -4.04 | < 0.001 |
| | Unit 2 | 0.164 | 1.020 | 0.16 | 0.872 |
| ***Sf-FAN2* vs *Sf-GSL1*** | Intercept | -0.002 | 1.637 | 0.00 | 0.999 |
| | Depth | -0.002 | 0.004 | -0.63 | 0.526 |
| | Unit 1 | -2.588 | 1.370 | -1.89 | 0.059 |
| | Unit 2 | 2.459 | 1.427 | 1.72 | 0.085 |
| ***Sf-GSL3* vs *Sf-GSL1*** | Intercept | 1.412 | 1.449 | 0.97 | 0.330 |
| | Depth | -0.007 | 0.003 | -2.79 | 0.005 |
| | Unit 1 | 0.084 | 1.251 | 0.07 | 0.946 |
| | Unit 2 | 3.078 | 1.408 | 2.19 | 0.029 |
| ***Sf-FAN1* vs *Sf-GSL1*** | Intercept | -0.205 | 1.277 | -0.161 | 0.872 |
| | Depth | 0.003 | 0.003 | 0.96 | 0.339 |
| | Unit 1 | -3.042 | 0.975 | -3.12 | 0.002 |
| | Unit 2 | 1.964 | 1.090 | 1.80 | 0.072 |

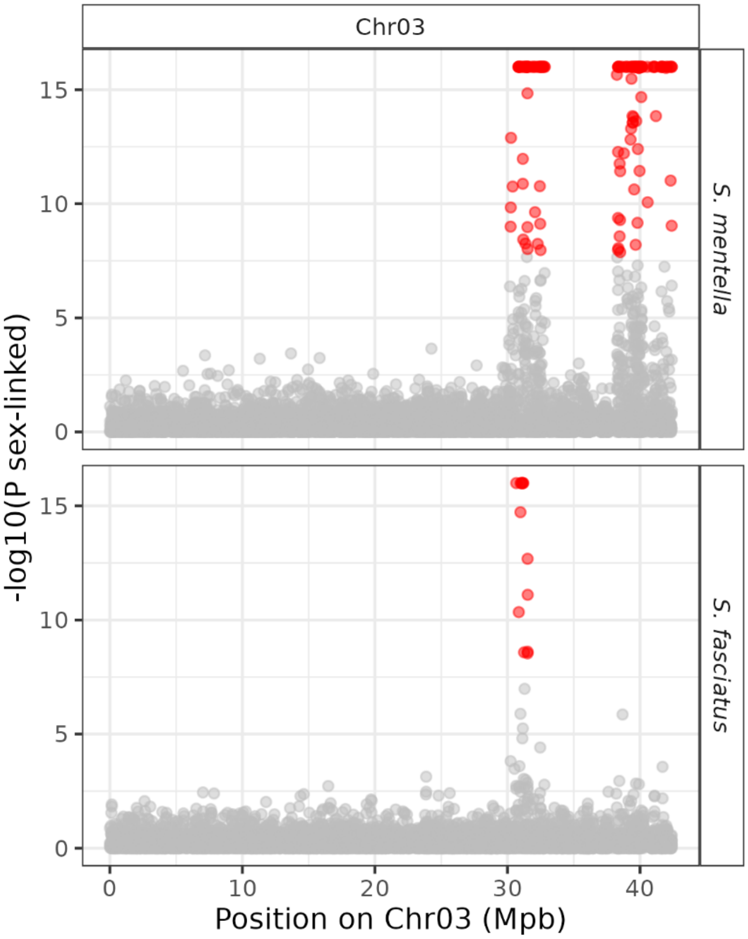


**Figure S1. RADsex results showing sex bias regions on chromosome 3 for *S. mentella* and *S. fasciatus*.** Each point represents a read, and red points indicates a probability of association with sex with a *p*-value < 0.05 after Bonferroni correction.

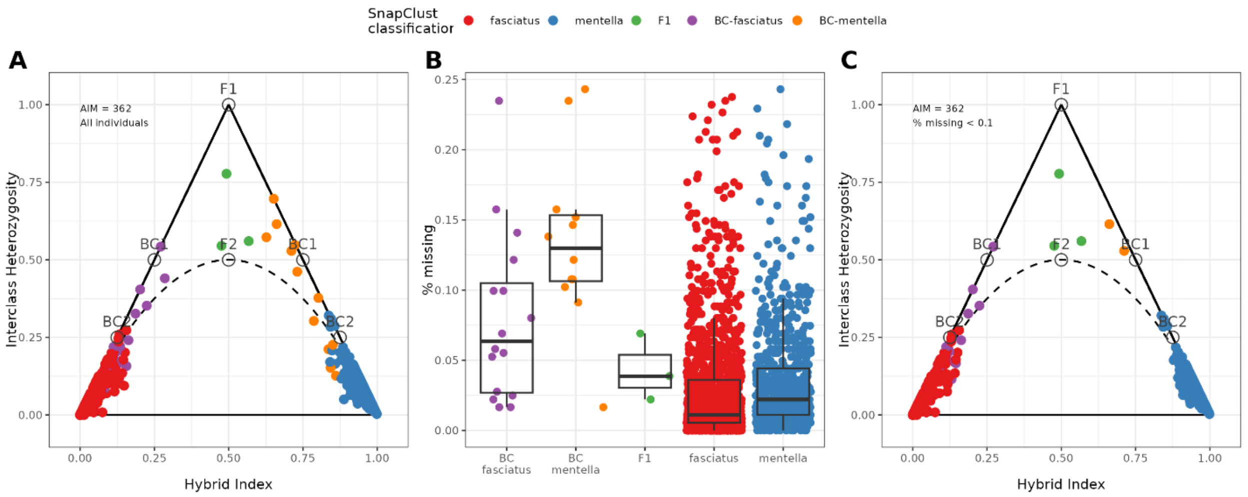


**Figure S2. Exploration with triangulaR of suspected hybrids and backcrossed individuals.** (A) Triangular plot using all individuals. (B) Percentage of missing values on the 362 AIM used by triangular. (C) Triangular plot using individuals with less than 10% missing values. Each point represents an individual, colored by it assigned category using SnapClust. Theoretical values are indicated in gray on the triangular plots.

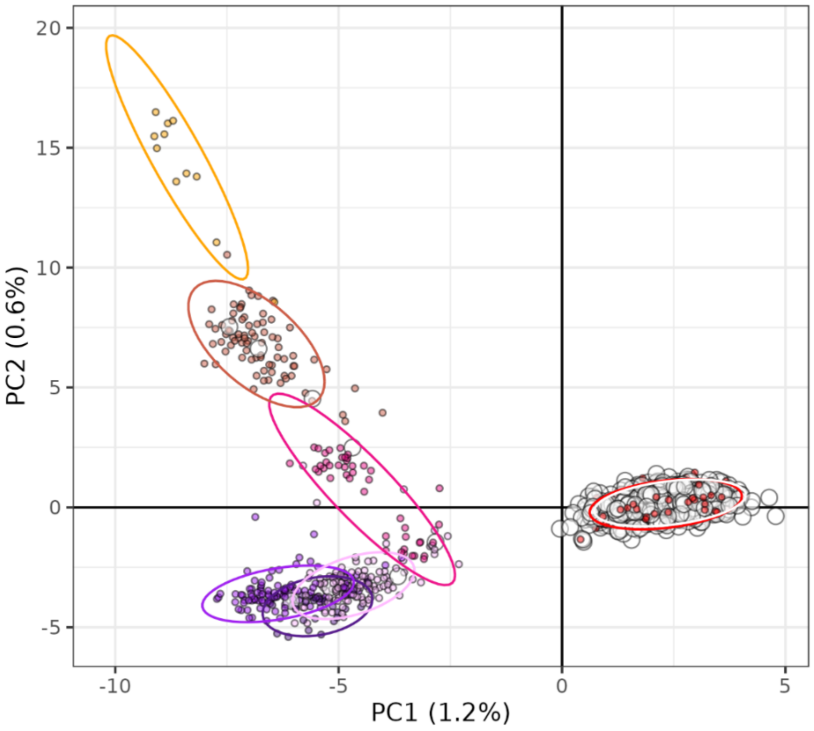


**Figure S3. Principal component analysis using all *S. fasciatus*, including the 795 small fish from the 2000s cohort sampled in 2005.** The 2000s cohort individuals were genotyped at the 16,333 SNPs of the *S. fasciatus* dataset and are represented by the large white dots. All other individuals are represented by small dots and colored following genetic group assignment by Snapclust at K = 7 (see Fig. 4A).

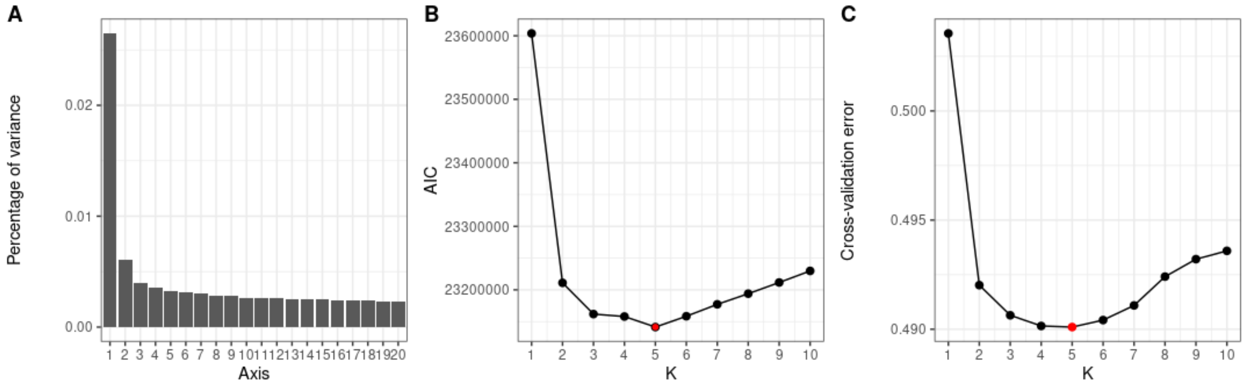


**Figure S4. Diagnostic plots to explore the structure in *S. mentella* and define the number of groups in the dataset.** (A) Percentage of variance explained by the 20 first axes of the PCA. (B) AIC for Snapclust clustering models from K = 1 to 10. Based on AIC value, K= 5 (red dot) was selected. (C) Cross-validation error of Admixture from k = 1 to 10. K = 5 (red dot) was the best value, but no further relevant structure was detected after K = 4. See Fig. S5 for results at K = 5.

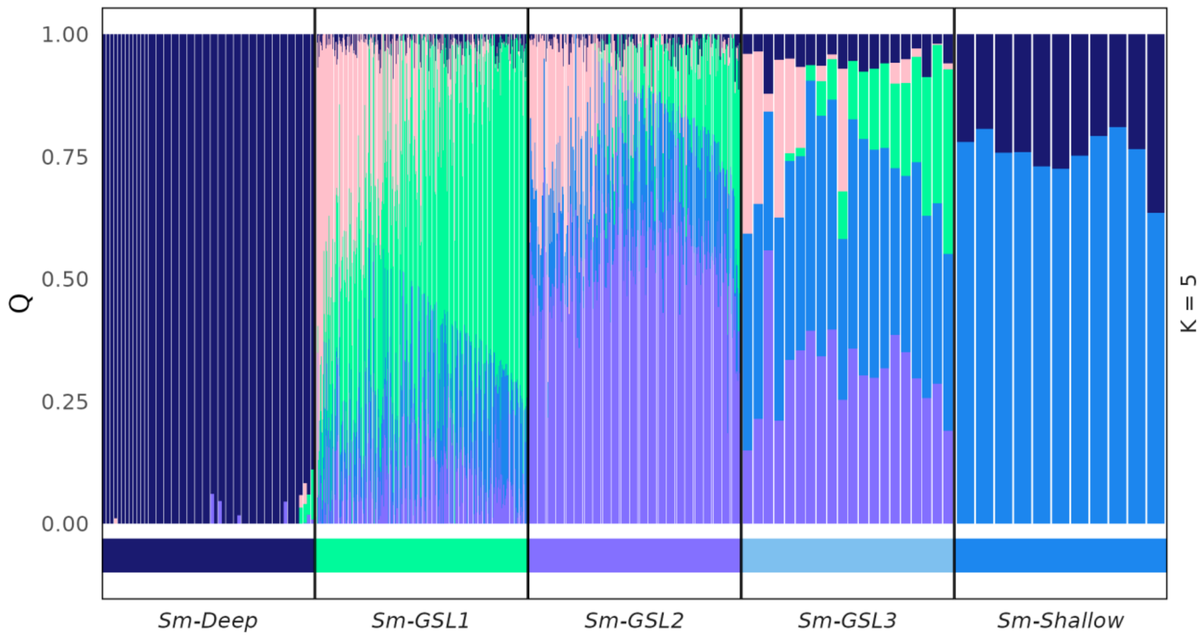


**Figure S5. Membership probability (Q) observed at Admixture K = 5 for *S. mentella.***

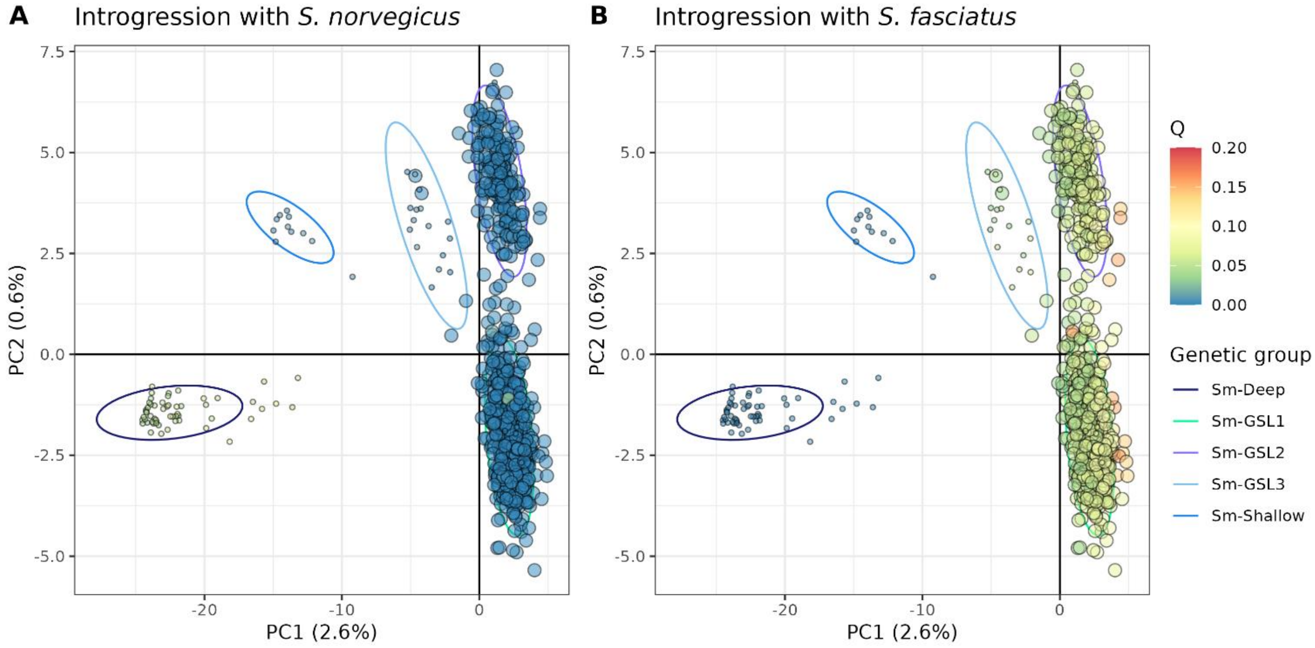


**Figure S6. PCA of *S. mentella* colored with membership probabilities (Q) of *Sebastes spp.* Admixture analysis to expose introgression with (A) *S. norvegicus* and (B) *S. fasciatus*.** Larger dots denoted recent sample and ellipses referred to the genetic groups assigned by SnapClust at K = 5.

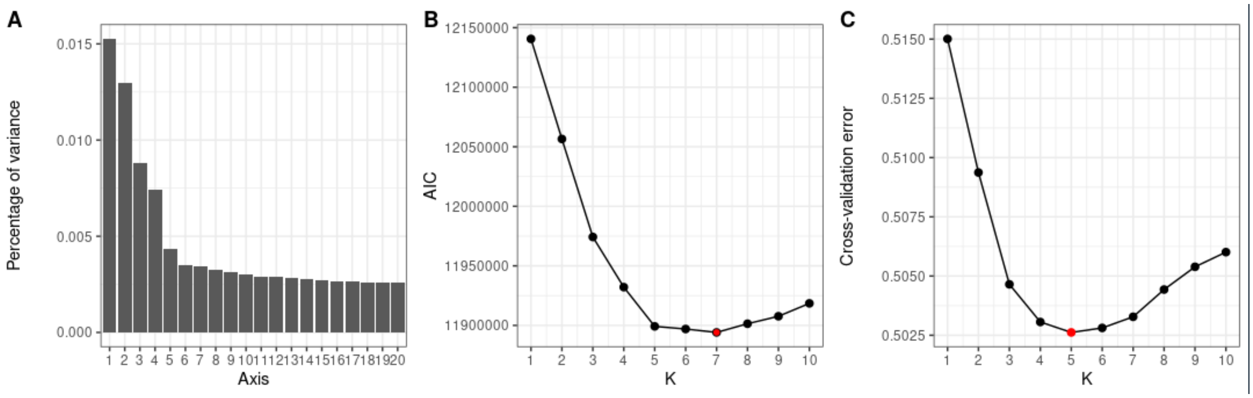


**Figure S7. Diagnostic plots to explore the structure in *S. fasciatus* and define the number of groups in the dataset.** (A) Percentage of variance explained by the 20 first axes of the PCA. (B) AIC for Snapclust clustering model from K = 1 to 10. Based on AIC value, K= 7 (red dot) was selected. (C) Cross-validation error of Admixture from k = 1 to 10. K = 5 (red dot) was the best value, but relevant structure was detected up to K = 7.

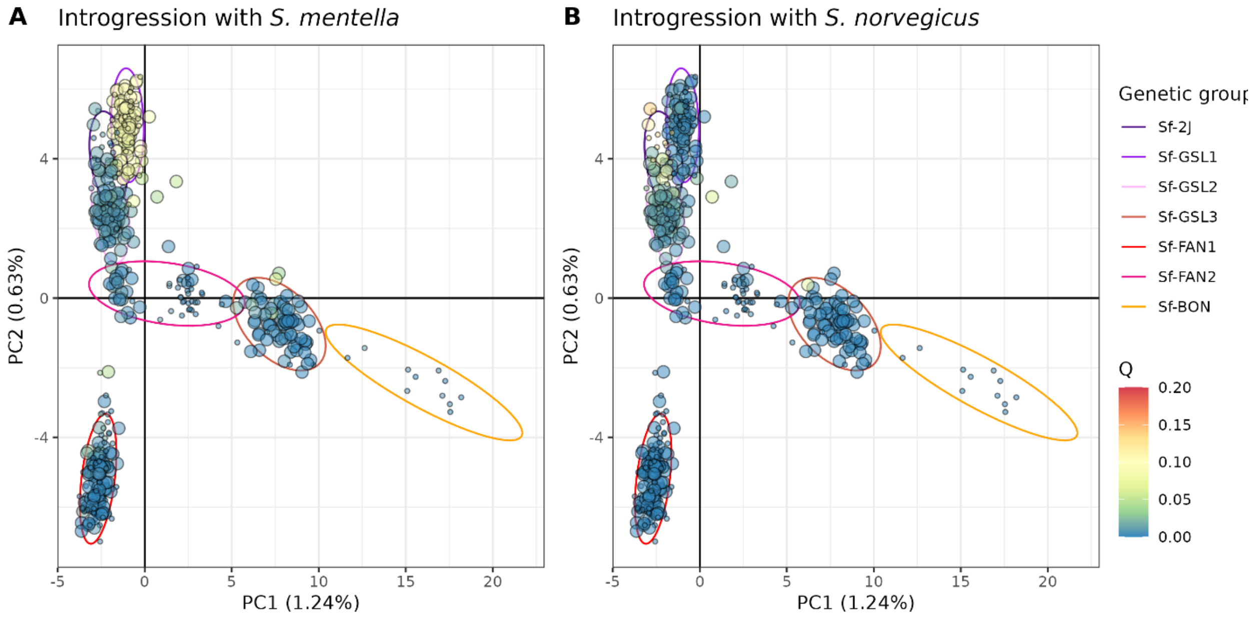


**Fig. S8. PCA of *S. fasciatus* colored with membership probabilities (Q) of *Sebastes spp.* Admixture analysis to expose introgression with (A) *S. mentella* and (B) *S. norvegicus*.** Larger dots denoted recent samples and ellipses referred to the genetic groups assigned by SnapClust at K = 7.
